# Copy number losses of oncogenes and gains of tumor suppressor genes generate common driver events of human cancer

**DOI:** 10.1101/2023.08.05.552104

**Authors:** Elizaveta Besedina, Fran Supek

## Abstract

Cancer driver genes can be under positive selection for various types of genetic alterations, including gain-of-function or loss-of-function point mutations (single-nucleotide variants, SNV), small indels, copy number alterations (CNA) and other structural variants. We studied the landscape of interactions between these different types of alterations affecting the same gene by a statistical method, MutMatch, which can test for significant differences in selection, while accounting for various causes of mutation risk heterogeneity. Analyzing ∼18,000 cancer exomes and genomes, we found that known oncogenes simultaneously exhibit signatures of positive selection and also negative selection, where the latter can mask the former. Consistently, focussing on known positively selected regions identifies additional tumor types where an oncogene is relevant. Next, we characterized the landscape of CNA-dependent selection effects, revealing a general trend of increased positive selection on oncogene mutations not only upon CNA gains but also upon CNA deletions. Conversely, we observe a positive interaction between mutations and CNA gains in tumor suppressor genes. Thus, two-hit events involving point mutations and CNA are universally observed on driver genes regardless of the type of CNA, and may signal new therapeutic opportunities that have been overlooked. An explicit focus on the somatic CNA two-hit events can identify additional driver genes relevant to a tumor type. By a global analysis of CNA-selection effects across many driver genes and tissues, we identified at least four independently varying signatures, and thus generated a comprehensive, data-driven classification of cancer genes by mechanisms of (in)activation by genetic alterations.

## INTRODUCTION

A deviation of observed somatic mutation rates compared to the expected baseline mutation rate provides evidence for selection during tumor evolution. The abundance of cancer exome and whole-genome sequences has accelerated the discovery of positively-selected cancer driver genes (and revealed an apparent rarity of negatively selected genes) by examining the local mutation rate of somatic single-nucleotide variants (SNVs) and indels and/or occurrence of hotspots thereof (1–5). However the sheer abundance of non-selected, passenger mutations compared to the few driver mutations makes it imperative to carefully account for the mutation rate baseline. This is a difficult task, bearing in mind the confounding by mutation risk heterogeneity, which occurs at various (overlapping) genomic scales, ranging from individual oligonucleotides over the kilobase gene-scale to megabase domain-scale variability in mutation risk (6–8). Various studies have addressed this challenge by modeling the mutation rate baseline based on covariates, such as DNA replication time, gene expression level, chromatin modifications, and the mutation type and the oligonucleotide (usually trinucleotide) context. Any significant deviation from the mutation rate modeled thusly is considered as a genomic signature of selection (1, 3, 5, 8, 9).

In addition to point mutations and small indels, also the somatic copy number alterations (CNAs) are commonly observed in cancer genomes and can drive tumorigenesis (10–12) by gene dosage changes and/or gene regulation changes. However, while some CNA are selected, as with the SNVs the CNAs can be very common and the majority of the genes affected by a CNA are non-selected passengers. Methods to identify these positively-selected driver CNA from genomic data are developing; they are based on recurrence of CNA events, sometimes incorporating external data from gene networks, multi-omic analyses or genetic screens (13–17). A particular difficulty with ascertaining selection on CNA is pinpointing which is the causal gene in a CNA segment, which often affects many neighboring genes. However, the cases where there is a simultaneous occurrence of point mutations at the CNA-affected locus may indicate the selection upon CNA in a particular gene. This is based on the fact that mutations and CNA can be epistatic in the same driver gene, reflecting diverse molecular mechanisms by which mutations and CNA activate/inactivate cancer genes.

An archetypal example of this are the SNVs or indels interacting with CNA deletions at the same locus, thus affecting two alleles of a tumor suppressor gene (TSG). This two-hit mechanism is common for deleterious germline variants in cancer predisposition genes, first discovered for *RB1*(18), and later shown for various other cancer risk genes including *ATM* and *BRCA1* (19). In addition to germline genes, two-hit inactivation mechanisms via CNA are broadly relevant also for somatic SNV mutations and can also involve epigenetic silencing of TSG allele (of note, the two-hits can also occur by a copy number-neutral loss-of-heterozygosity, which are not technically CNA) (20–23). Not only TSG, but also oncogenes (OG) were appreciated to be affected with selected CNA when the same locus bears point mutations. In particular, the dosage of the mutant allele tends to be increased relative to the *wild-type* allele (20, 23, 24). Such allelic imbalances in oncogenes were well studied experimentally for the *RAS* genes, prominently *KRAS* (25, 26), and genomics studies implicate at least several other oncogenes such as *KIT* and *EGFR* (20, 23).

In addition to the positive selection on TSG and OG, a point of special interest when considering interaction of SNVs and CNAs is negative somatic selection. Genomic signatures thereof in tumor genomes are very subtle (1, 2, 4), and so for most genes in most cancer types below threshold of detection. However, the (highly impactful but relatively infrequent) nonsense mutations do appear underrepresented in essential genes and oncogenes considered as a group (27–29), and there is some evidence that so are frameshifting indels (30). Considering selection on SNVs jointly with CNAs may clarify the signatures of negative selection, which was reported to be increased in hemizygous regions (1, 28, 31)., affecting for instance, the *POLR2A* essential gene (subunit of RNA polymerase II), genes encoding essential protein complex members as well as protein translation genes, and spliceosome genes (4, 20, 31). However, for most individual genes signatures of negative somatic selection remain elusive, and it is not clear whether somatic missense mutations are purged by selection in the CNA-neutral or CNA-gain chromosomal segments.

These prior studies reported prevalent interaction between somatic point mutations and CNA by measuring the statistical co-occurence of mutations and CNAs across tumors, or by studying the allelic imbalance of mutations in driver genes due to CNA. However the fitness effect of the altered mutant allele dosage remains understudied, since the rate of ‘neutral mutations was not systematically modeled analyses. This is particularly relevant for studies of driver CNA, because it is anticipated that occurrence CNAs — even if selectively neutral — may confound the local neutral mutation rate estimation. Baseline mutation rates are modeled per haploid gene/genome; an increased or decreased amount of DNA in a locus due to a CNA increases or decreases the observed mutation density per haploid gene. The state-of-the-art covariate-based methods for measuring selection (thus identifying driver genes) from point-mutation patterns do not consider CNA states and are thus naive to such confounding. While the classical evolutionary biology approach using dN/dS tests does inherently control for this (since the mutation rate baseline is located within the gene) however, because of low numbers of synonymous somatic mutations, the dN/dS test is less useful in practice, save for extremely large sets of cancer genomes/exomes (1, 4). Thus another approach that controls for local neutral mutation rate changes caused by CNAs is helpful for measuring interactions between selection on mutations and CNA, preventing false-positive and false-negative discoveries.

In this study, we apply a statistical method for cancer genome analysis (MutMatch) that estimates local mutation rates by a covariate-free method, which draws only on observed mutations in putatively neutral exonic regions matched by mutation rates to the gene-of-interest, and controls for effects of chromosome arm-level and segmental CNA that affect multiple genes. Using this, we rigorously characterize selection on genetic interactions between driver SNVs and CNA in the same cancer driver gene. This revealed that copy number losses were frequently associated with positive selection not only in TSG, as per known two-hit mechanism, but that CNA losses were also linked with positive selection in oncogenes. Conversely, copy number gains were associated with increased selection not only in oncogenes but also in TSG, increasing the dosage of the mutant allele in both cases. Next, we characterize signatures of selection in oncogenes upon stratifying by known positively-selected gene regions, and additionally by CNA states. This revealed that oncogenes are commonly under negative selection, including, remarkably, that they are under negative selection also in some tumor types where they are not commonly mutated. Further, factoring out negative selection increases sensitivity to find plausible long-tail driver events in oncogenes, such as *EGFR* driver mutations in luminal breast cancer and in head-and-neck cancer, and similarly so new drivers can be identified by simultaneous consideration of CNA and mutations. Finally, a global analysis of selection on SNVs in different copy-number states of oncogenes and of TSG across ∼18,000 tumor genomes provides a comprehensive, data-driven, mechanistic atlas of cancer genes categorized by the types and tissue spectrum of genetic alterations, including both SNV and CNA and their interactions.

## RESULTS

### A statistical method to test interaction between somatic selection on mutations and CNA in genes

We sought to rigorously measure somatic selection associated with a specific copy number state (neutral, gain or loss), which motivated us to develop a custom statistical genomic methodology. The MutMatch method compares the mutation rate in the coding exons of a gene of interest with the baseline mutation rate, estimated directly from observed mutations in matched loci that are presumably under neutral selection (**Fig. 1A**). By default, these matched loci are the coding regions of neighboring genes within 0.5 Mb both upstream and downstream, thus adjusting for the known domain-scale variability in somatic mutation rates, wherein such domains typically span multiple genes(3, 32, 33). Additionally, by using this approach, our MutMatch approach aims to control for confounding effects of larger segmental CNAs or arm-level CNA on neutral mutation rates, since the control genes are expected to be similarly affected by the CNA event as the gene-of-interest (central gene) (**Fig. 1A**). Additionally, the MutMatch method implements several refinements of the above-mentioned approach using neutral loci from neighboring gene exons. Firstly, controls on neighborhoods are performed by filtering genes in which neutral mutation rates were empirically found to deviate from the mutation rate within the rest of the neighborhood; see **Methods**. Importantly, we address the cases of CNA that are smaller than the neighborhood size by excluding the cases of neighboring genes in a different CNA state than the query (central) gene; this is considered separately for each tumor genome analyzed (described in Methods). Additionally, any trinucleotide composition differences between the query gene and the neighborhood genes are controlled for by a stratification into 96 mutational contexts, thus preventing confounding by differential activity of mutational signatures (**Methods**). Furthermore, to ensure that the method operates correctly on sparse data sets, we tested for biases during estimation on low mutation counts (**Supplementary Fig. S1A**) and implemented a randomization procedure that adjusts the observed effect sizes, based on a scrambled mutation counts baseline (**Methods**).

**Figure 1.**
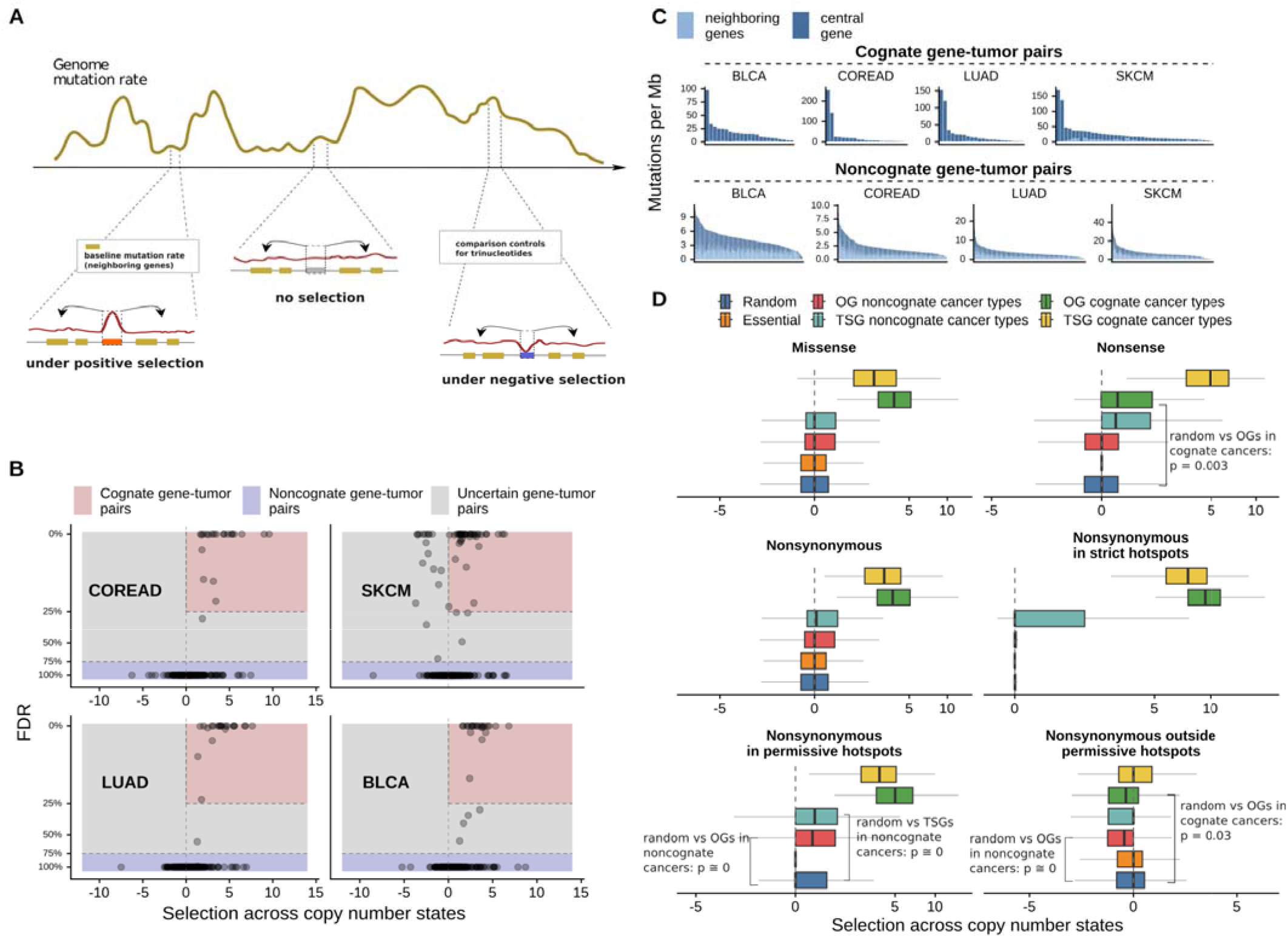
Estimates of somatic selection in various driver gene groups, within and outside the known positively selected hotspot loci. **A**. Schematic depiction of the MutMatch method to estimate somatic selection by using exonic regions of neighboring genes as a mutation rate baseline to compare against. **B.** Annotation of cognate and noncognate driver gene-tissue pairs is based on the selection estimates (log2 mutation rate fold-difference between the central gene and the baseline rate estimated from neighboring genes) and its significance level. **C.** Mutation rate in cancer genes and their corresponding neighboring genes for gene-tumor pairs defined as cognate and noncognate in four example cancer types (COREAD, SKCM, BLCA and LUAD). **D.** Strength of selection, here measured across all copy number states, estimated using neighboring genes as a mutational baseline. The brackets are used to highlight comparisons of interest, as commented in the Results section of the text. P-values are by Mann-Whitney U-test, two-tailed. The nonsynonymous mutations are the missense and the nonsense mutations considered together.

We applied the MutMatch method to aggregated mutational counts from various WES and WGS data sets, comprising over 18,000 tumor samples in total (see **Methods** for sources). We applied MutMatch separately for each cancer type and for every gene to obtain the selection effect estimates on mutations across copy number gain, loss and neutral states.

### Overview of analyses and cancer genomes and gene sets considered

Our first aim was to separately estimate selection on known gain-of-function (GoF) versus loss-of-function (LoF) mutations in coding exons of known oncogenes (OG), tumor suppressor genes (TSG), cell-essential genes (CEG2) (34), as well as a negative control set of random genes from which we excluded the OG, TSG and CEG (see **Methods** for curation of gene sets and data sources). To this end, we considered separately the nonsynonymous mutations in recurrent hotspots (the vast majority are missense), the truncating mutations (nonsense), and finally missense mutations outside of known hotspots. These three sets serve as representatives of GoF mutations, LoF mutations, and a mixed group of mutations that can be under varying degrees and types of selection, as investigated below.

In the initial analysis, we quantified signatures of selection across different gene sets –– random genes, OG, TSG, and CEG2 — and cancer types, considering the copy number neutral state (note that henceforth we call this neutral state “diploid” for simplicity, notwithstanding whole-genome duplication events). This was performed by including the CNA category (gain, diploid or loss) as a covariate in the regression model that additionally included an interaction term with this CNA variable; the interaction term would measure conditional selection upon CNA. In the initial analysis, the coefficients of the reference CNA state in the regression (this is the “diploid” i.e. copy number neutral state; see **Methods**) were studied. The interpretation of the regression coefficient here is the log fold-enrichment in relative mutation rate of a certain type of mutations in a gene, compared to that in the neutral control (neighboring) genes, after adjusting for gene length and trinucleotide composition.

For each cancer gene, we divided the cancer types into two categories (**Fig. 1B,C**): those where a gene was significantly positively selected (“cognate” cancer types for the gene) and those where it was not strongly selected (“noncognate” cancer types). This classification was performed using an operational definition based on positive selection estimates for mutations –– detected either on missense, or on nonsense, or on any nonsynonymous –– measured across all copy number states; see **Methods**. Cognate genes in a cancer type were defined as having FDR≤25% in that cancer type, while noncognate genes had FDR≥75% in that cancer type, and genes with FDR 25%-75% were considered uncertain and did not count towards either group (**Fig. 1B**). Our cognate/noncognate definitions broadly matched the cancer type-specificity spectrum from another recent comprehensive catalog derived via the MutPanning method (9) (**Supplementary Fig. S1B**).

As a control, the nonsense mutations in TSG in cognate cancer types were under a very strong positive selection (median = 4.9 (log_2_ fold-enrichment in mutation rates across all cognate gene-cancer type pairs, **Fig 1D**), while the TSG were under weaker positive selection in the non-cognate cancer types (median = 0.42, **Fig 1D**). We note that there is (modest) selection for nonsense mutations in some of the TSGs in the cancer types that we defined as noncognate (upper quartile = 1.59), implying that our following analyses that contrast cognate and noncognate genes are conservatively biased. Moreover, our classification of cognate and noncognate cancer types is independent from gene copy-numbers (the regression model did not include CNA information). As a side note, when considering OGs, there was a modest enrichment of nonsense mutations in some cognate cancer types for the OG (median was not notable at 0.48, however upper quartile = 1.66, **Fig 1D**), but not so in the non-cognate cancer types. This may be explained by either of the two (relatively rarely occurring) scenarios: instances of GoF truncating mutations in Ogs (35–37) and/or by genes that may act as either OG or TSG depending on the cancer type (38, 39) and that we classed as OGs.

### Signatures of selection in known hotspots identify additional cancer types where an oncogene is causal

Mutations in recurrent hotspots within coding regions of cancer driver genes are extremely likely to be causal, while the rest of mutations in cancer genes represent a mix of causal and other mutations in varied proportions. Next, we asked whether considering these two groups of mutations separately may increase power to identify moderate-effect cancer driver genes in smaller cohorts (e.g. in individual cancer types), and additionally whether it can clarify mechanisms of driver gene inactivation. For this analysis, CNA are not explicitly considered, instead we focus on selection pooled across all CNA states (**Fig 1D**; note that these analyses nonetheless control for possible confounding of CNA on baseline mutation rates). Additionally we also consider this analysis via selection acting only on the diploid state (**Supplementary Fig. S2A**). We considered two definitions of hotspot loci: (i) strict hotspots with likely functional effects as defined previously by Trevino (40) and available for n=283 cancer genes under consideration here; (ii) permissive hotspots detected in this study, based exclusively on recurrence of mutations and therefore possibly containing non-selected hotspots, but are available for a larger set of n=1240 genes and are derived from a newer, genomic dataset thus with likely better coverage of hotspots.

Expectedly, a much stronger positive selection was observed in strict hotspot sites than for all nonsynonymous mutations: for cognate OGs, median 9.34 *versus* 3.64 for selection across CNA states for all nonsynonymous mutation effects (**Fig. 1D).** Similarly in an additional analysis, we observe median 9.72 (log_2_ fold-enrichment in hotspots *versus* 3.22 considering selection in diploid state, **Supplementary Fig. S2A**). Also as expected, there was a more prominent signal in the strict hotspot set than in the permissive hotspot set; the latter did nonetheless exhibit stronger selection than the baseline (**Fig. 1D/Supplementary Fig. S2A**).

However interestingly, the selection in hotspots of OGs in the noncognate cancer types was significantly higher than in the control set of random genes (Mann-Whitney p = 0; median of OG hotspots log2 fold enrichment in noncognate cancer types 0.57, Q_3_ = 1.41, median of random genes ∼ 0, Q_3_ = 1.08). This suggests that some gene-tissue pairs from this noncognate (i.e. tentatively non-selected) group of OG-tissue pairs are in fact *bona fide* drivers in that cancer type. Presumably the selection signal for the OGs, evident within hotspots, was diluted by considering the entire gene locus when determining cognate and noncognate cancer types, meaning the OG did not pass our threshold of significance to be identified as a driver in a particular cancer type. Thus, driver genes can be identified better in various cancer types – including those where they are under weaker selection – by focussing the test for selection on the previously known hotspot-containing regions within a gene (**Supplementary Fig. S3)**.

Such apparently “noncognate” oncogenes with the strongest selection in the hotspots are shown in **Fig. 2C** (using the stringent hotspot definition), and here we highlight two interesting examples involving the *EGFR* and *ERBB2* genes. There is apparent selection on hotspot-located mutations in *EGFR* in the luminal subtypes of breast cancer (BRCA) (log_2_ fold-enrichment=5.72 across all three copy number states; p=8.3*10^-6^, FDR=2.9*10^-4^). *EGFR* is considered a lung adenocarcinoma and brain cancer driver gene (9, 41) and indeed in our analysis *EGFR* was identified as cognate only in GBM, LGG and LUAD (at FDR≤25%, **Fig. 2F)**. Given the crucial roles of the closely related gene *ERBB2* (*HER2*) in breast cancer, where *ERBB2* is often amplified and/or overexpressed, a role for *EGFR* alterations in breast cancer would not be unexpected. Indeed, this was considered for the triple-negative subtype (42, 43) and here we provide evidence that *EGFR* may also be a mutation-activated driver gene in the luminal breast cancer subtypes (**Fig. 2C-F)**. Next, the amplification-activated oncogene *ERBB2* was significant in this test, where we found selected hotspot point mutations in liver, uterine, kidney, melanoma and head-and-neck cancer (FDR≤25%, at **Fig. 2F**), while *ERBB2* was not marked as cognate in these cancer types (according to our annotation based on across-CNA states selection effects) nor according to MutPanning or Cancer Gene Census (CGC) annotations (9, 41).

**Figure 2.**
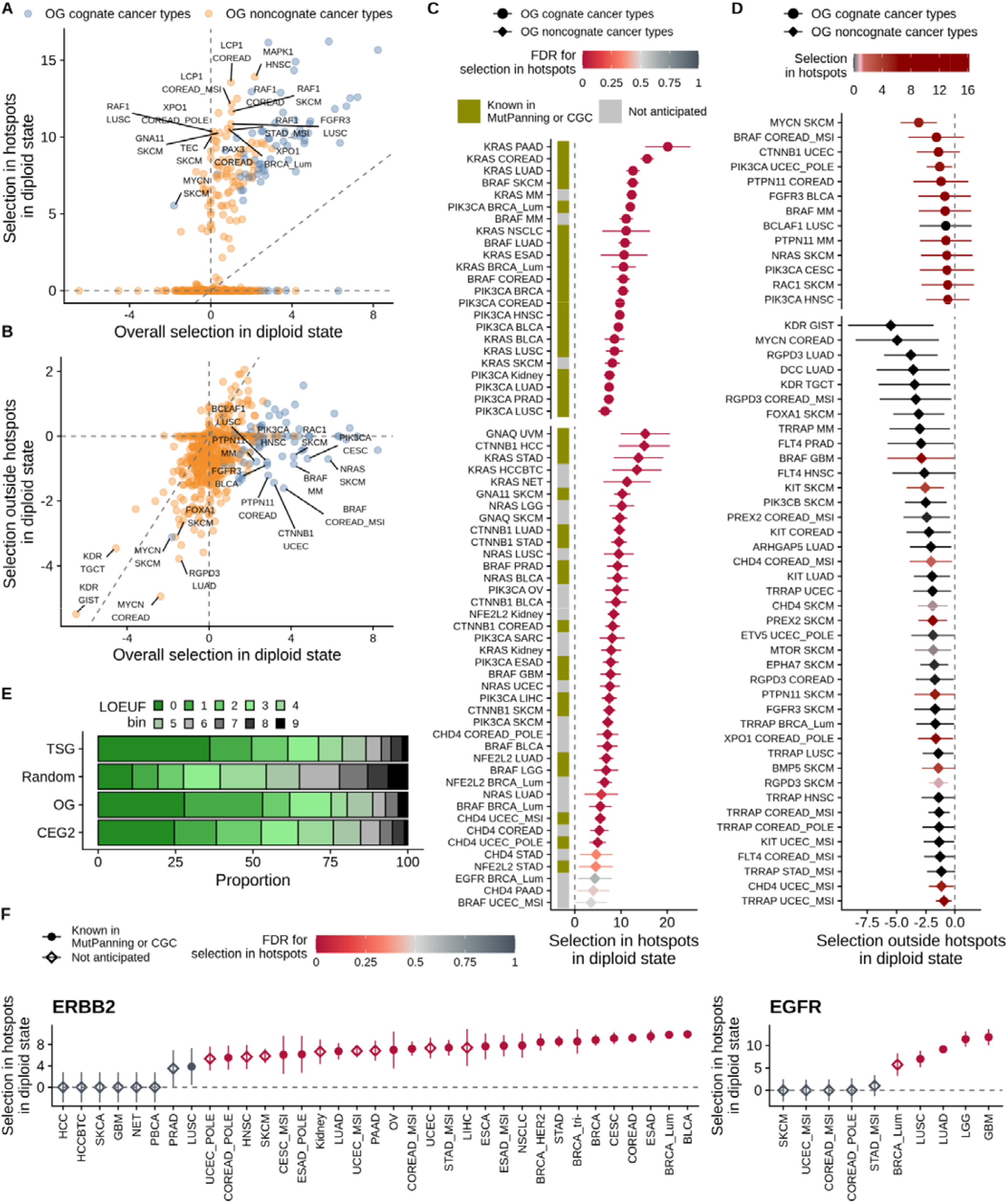
Positive selection and negative selection on oncogenes across various cancer types. **A**. Positive selection on oncogenes is observed within known hotspots (high values on y axis) in some cancer types that are annotated as noncognate (weak overall selection [x axis], orange points). One point corresponds to one gene-tumor type combination. **B.** Negative selection in oncogenes can often be observed after removing hotspot loci (y axis). This is seen both in the oncogenes that are overall under positive selection in a cancer type (cognate cancer types, blue points) and those that are not (noncognate, orange points). **C.** Oncogenes with the strongest positive selection effects in strict hotspots, here showing the top-50 frequently mutated MutPanning genes. Top, examples of cognate genes as positive controls, bottom, all significant noncognate genes. The color of the points represents the FDR for selection in hotspots, color of the left bar encodes whether a gene-tumor type combination is a known driver according to the MutPanning of CGC lists. **D**. Oncogenes with the strongest negative selection effects outside permissive hotspots. Error bars show 95% CI. Grey points correspond to the genes lacking positive selection in strict hotspots but under negative selection (“noncanonical oncogene addiction”), red points show gene-tissue pairs with opposing positive and negative selection inside and outside of hotspots, respectively. Genes with the strongest negative selection outside of permissive hotspots are shown. **E.** LOEUF scores distribution measuring population genomic constraint from Loss-of-Function (LoF) variants, with lower scores correlating with haploinsufficiency in the human germline. **F.** Positive selection of point mutations in *ERBB2* and *EGFR* genes observed hotspots in different cancer types, either where it is a known driver (according to MutPanning or CGC) or not known.

Further examples of genes where signal of positive selection was clarified in hotspots include *CTNNB1*, *CHD4*, *XPO1*, *LCP1*, *FGFR3*, and *RAF1* in various cancer types (**Fig. 2C** showing a selection of major OGs in noncognate cancer types, prominently *BRAF, KRAS, NRAS, CTNNB1* and *PIK3CA*; full complement of genes shown in **Supplementary Fig. S3B)**. Overall, 86 OG-cancer type pairs were significant in the hotspot-enrichment analysis (at FDR≤25%) but far from significant (FDR≥75% i.e. “noncognate”) in the general enrichment of mutations. Similarly, 91 TSG-cancer type pairs were significant in the hotspot test only (FDR≤25%) but not in the general enrichment analysis (FDR≥75%). Thus, focussing on hotspot regions enriches for positive selection signal, allowing to identify with higher statistical power which driver genes are relevant to which cancer types. Of note, this is related to but distinct from methods that search for significant clustering of mutations, thus defining hotspots *de novo* and identifying new cancer genes (44–46). The approach described herein relies on known hotspots and thus is not able to identify new cancer genes, but is able to measure the cancer type spectrum of known genes with better signal-to-noise.

### Prevalent signatures of negative selection on nonsynonymous mutations in oncogenes

Considering the diploid state, a suggestive signal of negative selection was found on nonsense mutations in some OGs: the lower quartile of the nonsense mutation fitness effects was trending towards more negative values in OGs, compared to random genes ((log_2_ mutation rate fold-enrichment = –0.49 in OG in noncognate cancer types and –0.01 in random genes, p=0.099 by permutation test, **Supplementary Fig. S2A**; we note the median was at 0 thus the majority of OG do not show a signal in this test, and neither did essential genes). This is in line with recent reports, where purifying selection for nonsense somatic mutations was shown on OGs in a group analysis (27, 29). While we did not identify significant individual genes under negative selection, probably due to statistical power issues, some top-ranked candidates were *KMT2A* and *XPO1* in COREAD-POLE mutant cancer (FDR=20% and 37%, respectively), in *CDH17* in in UCEC-POLE mutant cancer (FDR=59%), and *IL6ST* in BRCA-Luminal subtype (FDR=60%) (others in **Supplementary Fig. S2B**). Similarly, among the known cell-essential genes in the CEG2 set, we do not observe a notable footprint of negative selection on nonsense mutations nor on all nonsynonymous mutations (neither in the diploid state, nor when considering all three copy number states jointly), consistent with prior reports (1, 28). However, selection on nonsynonymous mutations were significantly lower for the upper quartile (Q_3_ of essential genes = 0.4, Q_3_ of random genes = 0.47, p ∼= 0 by permutation test), suggesting a subset of essential genes has detectable negative selection in cancer. We do recover the previously-reported *POLR2A* gene (31) having considerable (albeit nonsignificant) depletion of nonsynonymous mutation rates across three CNA states e.g. log2 fold enrichment of –1.32 in UCEC-POLE cancer type, –0.8 in BRCA-Lum, –0.79 in BRCA and LGG cancer types.

Motivated by the trend depletion in nonsense mutations in OGs, we next asked if the negative selection in OGs is seen in the (far more numerous) missense mutations. We hypothesized that the pattern of mutations on OGs might be explained not only by positive selection acting on GoF mutations, but also by a negative selection that acts to purge deleterious missense mutations in regions located outside of hotspots, where most GoF mutations are contained. To examine this, we estimated selection in hotspot-free gene regions, using a permissive set of hotspot loci for exclusion (see **Methods**) thus minimizing the effects of positive selection in the remaining regions (**Fig. 1D**, **Fig 2B)**.

Indeed, the out-of-hotspot nonsynonymous mutations in OGs in noncognate cancer types were selected negatively (**Fig. 1D**, first quartile of distribution for OGs Q_1_ = −0.84 and median = −0.29) in comparison to the set of random genes (in first quartile Q_1_ = −0.54 and median ∼0); this difference in medians is significant (p∼0 by Mann-Whitney test). Similarly, for the OGs in cognate cancer types, a shift towards negative selection from random genes was observed outside of hotspots (for OGs, Q_1_ = −0.81 and median = –0.24); difference to random genes p=0.028 by Mann-Whitney test). In other words, we estimate that OGs have ∼18% fewer mutations than expected in cognate cancer types and ∼22% fewer mutations than expected in noncognate cancer types (based on the median of the distribution, and normalizing by the random genes group). These estimates are conservative, as some of the subtly positively selected mutations may not have been removed together with the hotspots.

Our analyses indicate that there is negative selection on mutations in oncogenes after excluding the known GoF mutations in the positively-selected hotspots (**Fig. 2B)**. Therefore in the remainder of OG coding regions, some sites are essential for the activity of the gene and the oncogene itself is essential for the tumor.

### Noncanonical oncogene addiction, where oncogenes exhibit only negative but not positive selection signatures

Next, we asked if the signatures of negative selection seen on nonsynonymous mutations in oncogenes are contained to oncogenes that are positively selected in a given cancer type. We compared signals of selection on OGs inside strict hotspots (from Trevino et al.(40)), presumably under clear positive selection, against hotspot-free areas, here excluding the permissive set of hotspots, to minimize the signal of positive selection on this remainder of gene coding sequence (**Fig. 2A-B**; this analysis can consider the n=67 OGs that had available the definition of strict hotspots (40)). The oncogenes formed two clusters as expected: first, where hotspot-mutations were not strongly selected, which contained predominantly the genes from noncognate cancer types, and the second group that had a strong selection in hotspots and consisted mainly of cognate cancer types meaning that OGs were overall under positive selection.

Notably, the negatively selected oncogenes were similarly prevalent in both groups. We estimated that 43% and 50.6% of gene-tissue pairs were negatively selected (here defined as (log_2_ mutation rate fold-enrichment < –0.2 regardless of significance) in cognate and in noncognate cancer types, respectively. This suggests two possible mechanisms underlying selective pressures that remove deleterious mutations from OGs. The first mechanism acts on driver oncogenes (that are mostly in cognate cancer types) and prevents that the oncogene bearing a GoF mutation is inactivated by another deleterious mutation. This LoF mutation would reverse the fitness gain of the GoF mutation, and in addition may potentially incur an additional fitness loss because of “oncogene addiction” where the cell state after the GoF mutation is such that a withdrawal of its activity can be lethal. The second mechanism of negative selection on OGs highlighted here, however, demonstrates that tumors depend on the oncogene function even in some cases where that specific oncogene is not a driver of tumorigenesis in that particular cancer type. Herein we term this phenomenon “non-canonical oncogene addiction”, where a cancer cell is dependent on the function of an oncogene even without its activation by a GoF function being selected in that cancer type. For instance *MYCN*, *MTOR*, *BRAF*, *KIT*, *KDR* and *XPO1* genes are suggestive examples of genes from this group showing negative selection signals in apparently noncognate cancer types (**Fig. 2D**) with the effect sizes (log_2_ relative mutation risk depletion compared to baseline) ranging from –5.49 to –2.52 for top-10 hits, with significant FDRs ≤25% for some of them (*KDR* in GIST, *RGPD3* in LUAD, *KDR* in TGCT).

There is another intriguing implication of these signals of negative selection on OGs in the noncognate cancer types (i.e. where overall signal selection was not strongly positive in our analyses). Considered together with the subtle signal of positive selection in hotspots of these putatively noncognate OGs (see above) suggested that a fraction of these cancer types might have been wrongly annotated as noncognate in our (and likely other previous) analyses, because of confounding between positive and negative selection. In other words, negative selection may offset some part of the signal of positive selection on the same oncogene, reducing the power to detect the positive selection, causing a false-negative i.e. failure to identify this as a driver cancer gene in a given tumor type.

### Essential genes show modest signatures of negative selection in somatic cells

In addition to oncogenes, we further considered a set of known cell essential genes (CEG2 set, derived from cell line genetic screening data (34)), analyzed selection in diploid state on nonsynonymous mutations in a pan-cancer analysis across 13 major cancer types (those with the largest numbers of mutations in our data: BLCA, BRCA-Lum, COREAD, ESAD, HNSC, Kidney, LGG, LUAD, LUSC, MM, PAAD, PRAD and SKCM). Our results show that essential genes are indeed negatively selected in the pan-cancer analysis in diploid state (Mann-Whitney p =7.3*10^-4^, median of random genes = –0.06 and median of CEG2 genes = –0.15; for comparison, median of all OGs = 0.11 and for TSG = 0.14 in this test that does not distinguish cognate and noncognate cancer types for OG and TSG). Regarding establishing tissue-specificity of these negative selection signatures, given the overall subtle effect sizes, there was not enough power in the dataset to detect the significance in each cancer type separately. The known essentiality metrics correlated only very modestly with our negative selection estimates across the cancer genomes: CERES score (gene essentiality across cultured cancer cell lines, determined in CRISPR screening experiments (47)) at R=0.06 and LOUEF population genomic score at R=-0.07 **(Supplementary Fig. S4A-B**). Nonetheless, this CERES score (34) was more correlated with our estimates of negative selection in cancer, than was the LOEUF score prioritizing genes with human population constraint on germline variants (48) (measuring a dearth of germline LoF variants over an expectation; of note the CERES and LOEUF scores only weakly correlated at R = 0.16, **Supplementary Fig. S4D**), thus supporting the plausibility of the negative selection signal in tumor genomes. Overall, our results suggest that signatures of negative selection in cancer on point mutations are extant but very subtle, and do not currently allow prioritizing individual genes by essentiality in different cancer types.

On a related note, we also compared the distributions of CERES (cancer cell line genetic screening) essentiality scores in different groups of cancer genes, contrasting with random genes (**Fig. 2E, Supplementary Fig. S4E**). Cancer genes were more essential by this criterion (**Supplementary Fig. S4E**): Q_1_ = −0.19 and median = 0.22 for TSGs, and Q_1_ = −0.06 and median = 0.26 for OGs, compared to Q_1_=0.09 and median=0.32 for the random genes). A similar result was seen with the population-level LOEUF score (**Fig. 2E**): 64% and 65% of OGs and TSGs, respectively, had the LOEUF scores from the lowest 3 deciles (thus the expectation at random would be 30%). This suggests that functions of cancer genes, considered as a set, tend to be more essential at the level of the cell (CERES) and also at the organismal level (LOEUF). This result is consistent with the observed negative selection on mutations in oncogenes in cancer genomes, as we reported above, and suggests an intriguing possibility that some TSGs could be essential in cancer cells.

### Selection change upon somatic CNAs identifies additional cancer types where a gene acts as a driver

To evaluate how somatic CNAs affect the selection of somatic point mutations in genes, we performed two analyses. First, we estimated conditional selection on point mutations associated with a gene loss (i.e. a change in selection strength that is observed in tumor genomes where a gene copy is deleted, compared to copy-number neutral, **Fig. 3A**). Second, we estimated selection change upon a gene copy-gain (**Fig. 3A**).

**Figure 3:**
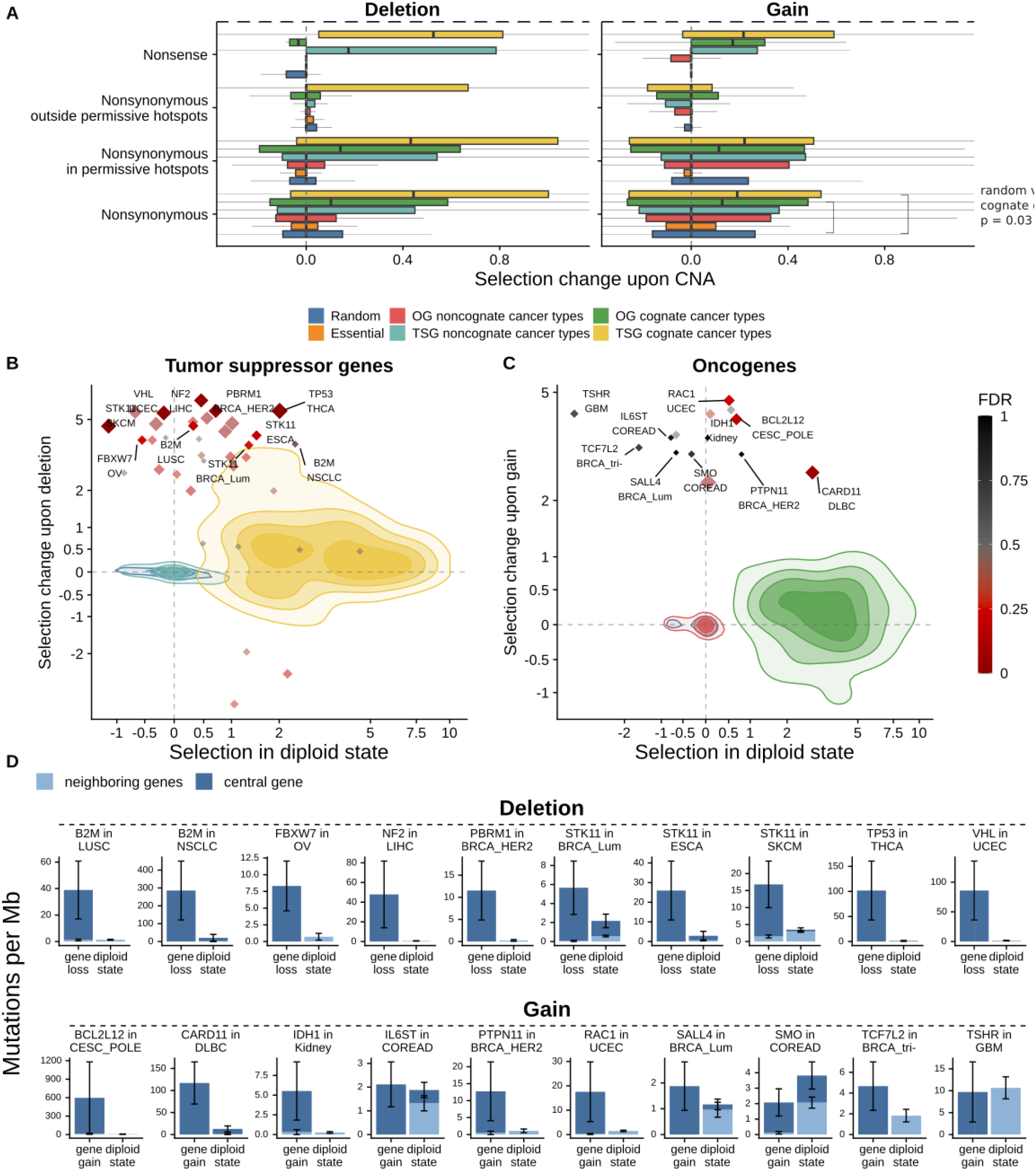
Positive interaction between mutation rates and CNA helps to identify new driving genes for cancer types. **A**. The change of selection strength between samples where genes are in the diploid state and where a gene copy was lost or gained, estimated using neighboring genes as a mutational baseline. **B.** Effect sizes of selection change upon deletion of random genes, TSGs in cognate and noncognate cancer types compared to the selection strength in diploid state. Colors encode gene groups as in **A**. Only the cancer types with the largest number of samples are shown; gene-tumor pairs with the lowest FDR of selection change upon deletion are labelled as individual points, while the rest of the gene-tumor type pairs are shown as a distribution (density plot). Axes are log-transformed with a smooth transition to linear scale around 0 (pseudo-log transformation (49)). **C.** Effect sizes of selection change upon gain of random genes, OGs in cognate and noncognate cancer types compared to the selection strength in diploid state. Visualization as in panel ***B***. **D.** Examples of newly-identified gene-tumor pairs, exhibiting increased mutation rate upon CNA loss or gain, compared to the neutral mutation rate in neighboring genes. Mutation frequencies (number of mutations per Mb per sample) shown in the plot were not adjusted for trinucleotide composition. Error bars are standard errors.

First, we checked for genomic evidence supporting the known two-hit model of TSG inactivation (18, 21). In agreement with this, selected nonsense mutations and deletions were co-occurring more than expected by chance in TSGs. (**Fig 3A**, median effect size for TSG nonsense mutations in diploid state = 4.62, while in loci with CNA deletions = 5.14 (log_2_ fold-enrichment in mutation risk). Interestingly, although the effect was strong for cognate cancer types, a distribution tail of positive interaction between selected nonsense mutations and deletions was also shown for TSGs in apparently non-cognate cancer types (we identified cognate versus non-cognate cancer types using selection estimates across all CNA states). In particular, the TSGs *NF2* in LIHC, *VHL* in UCEC, *B2M* in LUSC and NSCLC, *STK11* in SKCM and ESCA (FDR≤25%) and other noncognate gene-tumor pairs were not showing significant selection in diploid state but did show selection in tumors with deletion in the same gene (**Fig. 3B,D, Supplementary Fig. S5A**). This suggests that considering somatic mutations specifically in the genome segments harboring CNA deletions –– thus focussing on two-hit events –– has the potential to identify new driver genes for particular cancer types by enriching for causal mutations. This is by analogy with a germline variation analysis that monitored co-occurrence of pathogenic germline variants with somatic LOH to identify heritable cancer predisposition genes from genomic data (19). We estimated the number of gene-cancer type pairs that appeared to be under a positive selection change upon copy number events (here defined as: lower boundary of 95% CI of log_2_ mutation rate enrichment interaction term > 0.2). We found that 49 noncognate TSG-tumor type combinations were positively selected in tumor samples with deletion, suggesting they are bona fide TSGs in that tissue. For comparison, 193 TSG-tumor type combinations were selected (cognate) according to our definition across copy number states.

Second, by analogy to the above, we considered the case of OGs and how CNA data might be used to boost the power to identify driver oncogenes pertinent to certain cancer types. The two-hit pattern involving CNA gains was reported to be common in some oncogenes, not only in the well-known cases of highly amplified OGs such as *EGFR,* but also in the case of low-level gains affecting various genes (20, 23). Indeed, also in our data we find that in the CNA gain state, overall positive selection (on all nonsynonymous mutations) in OGs is modestly increased compared to the diploid state (median effect size for cognate OGs = 3.34 in gain versus 3.22 in diploid; **Fig 3A** shows interaction terms from the regression, which are the difference of the two effect sizes). Motivated by this, we hypothesized that by considering the CNA gain state, we may identify some driver genes selected in certain tissues that would otherwise be classified as non-cognate i.e. not significantly selected by our standard classification. Indeed at least some of the non-cognate OGs do show an increased selection in the permissive hotspots upon CNA gains (Q_3_ of the interaction term = 0.4, compared to Q_3_ of random genes set = 0.23, p=∼0 by permutation test; **Fig. 3A**; we note the medians are the same, meaning this principle would not apply to the majority of OGs). Indeed in some cases of apparently non-cognate OG-tumor type combinations, there was selection in the copy-number gained state (*BCL2L12* in CESC_POLE, *RAC1* in UCEC, *HMGA1* in BLCA and other gene-tumor pairs with FDR ≤25%) **(Fig. 3C-D, Supplementary Fig. S5B**). We found that 17 noncognate OG-tumor pairs were selected positively upon gene gain (with lower boundary of 95% CI of (log_2_ mutation rate enrichment > 0.2), compared to 134 OG-tumor type combinations found selected according to our definition across copy number states.

Finally, we further considered the case of negative selection on mutations in OGs, as we noted above, here hypothesizing additionally an interaction thereof with CNA state. This is motivated by an analogy with previous reports that essential genes in cancer may show signatures of negative selection specifically in hemizygous segments (1, 31). Indeed, here we also observed a similar pattern with OGs. There was a modest depletion in the rate of nonsense mutations in OGs upon deletion (negative conditional selection), compared to the baseline (median effect size of the interaction term from the model =-0.03 for OGs in cognate cancer types compared to ∼0 for random genes, **Fig. 3A**). This, however, was not significant, likely due to the small sample size, stemming from the relative rarity of nonsense mutations in oncogenes. Additionally, stronger negative selection was found inside permissive hotspots of OGs in the CNA-deleted state (interaction term: (log_2_ fold decrease in mutation rates upon deletion Q_1_ = –0.19 *versus* –0.07 for random genes, p = 0.0038 by randomization test). This indicates negative selection in hemizygous regions of some oncogenes. Therefore oncogenes, similarly as essential genes, appear to be under additionally increased negative selection in chromosomal segments that have undergone copy number losses.

### Positive selection on mutations is increased by deletions in OGs and gains in TSG

In addition to the known case of two-hit interactions between mutations and CNA gains in OGs, and mutations in CNA losses in TSGs, we hypothesized that the opposite case may also be selected. Recently it was reported that allele imbalance of mutations at OGs can result from deletions (20) and we thus turned to examine differential selection on point mutations in OGs upon deletions. For symmetry, we additionally checked the case of TSGs and amplifications.

Both in TSGs as in OGs, selection on all nonsynonymous mutations was stronger in samples with a CNA loss of the same cancer drive gene, comparing to a control set of genes (**Fig. 3A**) (effect size in the following text refers to an interaction term from the model, i.e. difference of (log_2_ mutation rate fold-enrichment, comparing the diploid state *versus* the CNA loss state). In particular, the differential selection effect size for random genes has the distribution upper quartile Q_3_ = 0.15 and median = 0, while for cognate OGs Q_3_ = 0.58 and median = 0.10 and for cognate TSGs Q_3_ = 1.00 and the median = 0.44 (**Fig. 3A**). Therefore the two-hit deletion effect in potentiating positive selection in at least some OGs (Q_3_: top 25% OG-cancer type combinations) is higher than the two-hit deletion effect on the median of cognate TSGs (**Fig. 3A**, **Fig. 4B-C)**). Next, as a validation we hypothesized that this two-hit OG deletion effect should become more pronounced in the case of mutations in hotspots, which are in OGs very strongly enriched with positively selected mutations. Indeed, supporting our observations, by considering the permissive hotspots Q_3_ = 0.63 for OGs and Q_3_ = 1.04 for TSGs, while for random genes the Q_3_ is expectedly much lower (Q_3_ = 0.04). Therefore, deletions in OGs — similarly as was known for TSGs — can in fact commonly increase tumor fitness via two-hit events.

**Figure 4.**
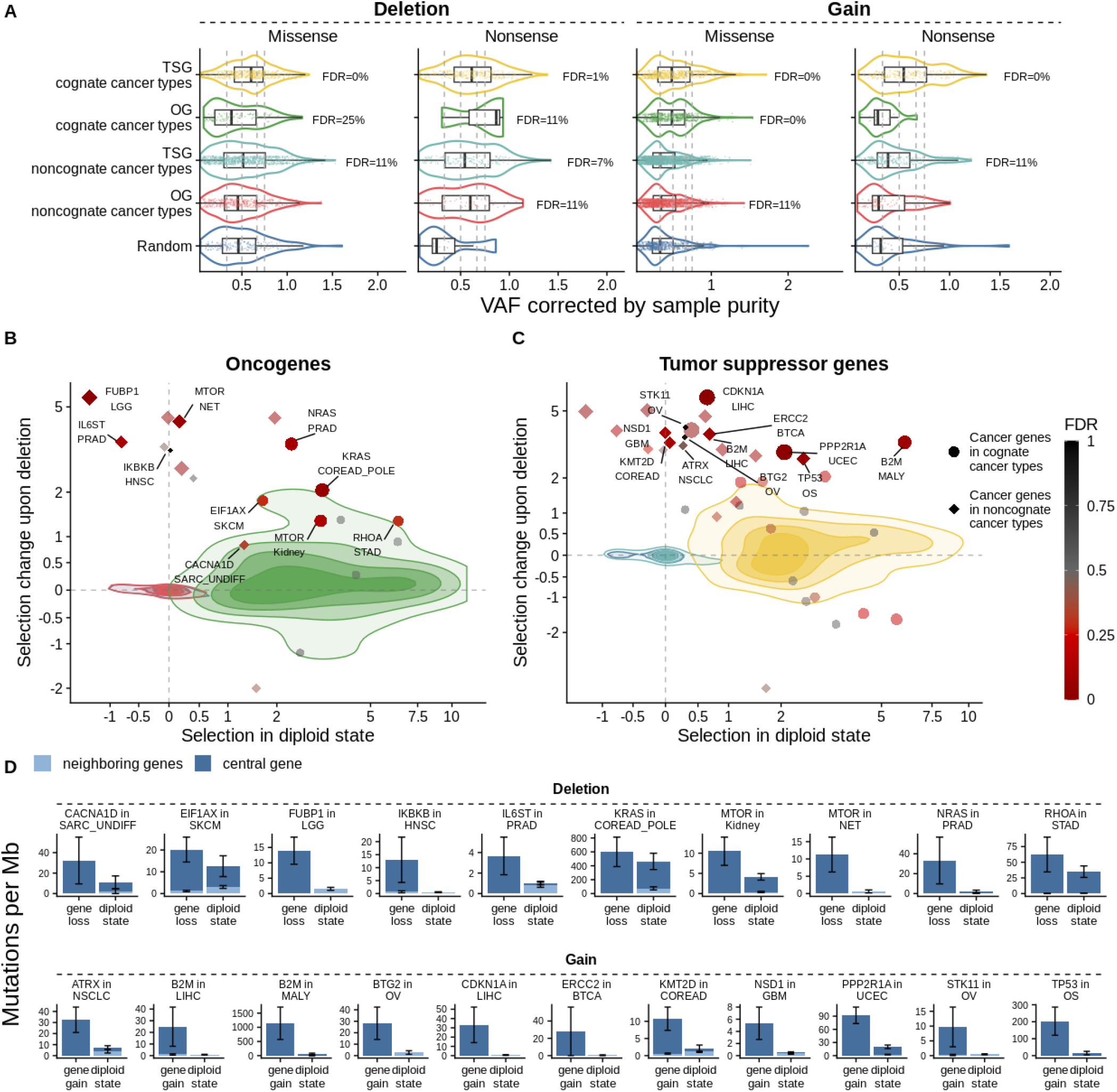
Prevalent association between CNA gains and positive selection in TSGs and CNA deletions and positive selection in OG. **A**. Mutation frequencies in samples with a gene loss or gene gain. One data point corresponds to one adjusted frequency of one missense or nonsense mutation for each tumor sample from TCGA. Adjusted p-values (Mann-Whitney test) are shown for FDR≤25%. **B.** Effect sizes of selection change upon deletion of random genes, OGs in cognate and noncognate cancer types compared to the selection strength in diploid state. Colors encode gene groups as in **A**. Only the cancer types with the largest number of samples are shown; gene-tumor pairs with the lowest FDR of selection change upon deletion are labelled as individual points, while the rest of the gene-tumor type pairs are shown as a distribution (density plot). Axes are log-transformed with a smooth transition to linear scale around 0 (49). **C.** Effect sizes of selection change upon gain of random genes, TSGs in cognate and noncognate cancer types compared to the selection strength in diploid state. Visualization as in panel ***B***. **D.** Examples of newly-identified gene-tumor pairs with increased mutation rate upon gene loss or gene gain compared to the neutral mutation rate in neighboring genes. Mutation frequencies (number of mutations per Mb per sample) shown in the plot were not adjusted for trinucleotide composition. Error bars are standard errors.

Next, we turn to consider how the selection of mutations in OG and TSG changes in tumor samples with a gene gain (**Fig. 3A**; effect sizes shown are differences of log_2_ mutation rate fold-enrichment, comparing diploid state and CNA gain state). We observed that driver missense mutations in OGs had, expectedly, an increased selection in samples with CNA gain in OGs (20, 23), compared to non-selected gene-tissue pairs (meaning: either the control set of random genes, or cancer genes in noncognate cancer types). However we observed that TSGs are also under increased selection upon gains, considering nonsense and missense mutations as potential drivers. In particular, the differential selection effect sizes (log_2_ fold increase in mutation rates upon gain, obtained from the interaction term in regression) for all nonsynonymous mutations was median = 0.19 and Q3 = 0.54 in cognate TSGs. For comparison, in noncognate TSG median = 0 and Q_3_ = 0.36 and in control random genes was median = 0 and Q3 = 0.26 (**Fig. 3A**). Remarkably, this difference in selection on TSGs upon CNA gains was slightly greater compared to the known two-hit effect on selection on OGs upon CNA gain (median = 0.13, Q_3_=0.48). Some of the tumor suppressor genes with an increase of selection upon CNA gain included *NSD1* in GBM (FDR=14%), *ERCC2* in BTCA (FDR=8%), and *STK11* in OV and *B2M* in LIHC (with non-significant FDRs) and others (listed in **Fig. 4C, Supplementary Fig. S6B**).

Overall, there is increased selection of mutations in OGs upon CNA deletions, suggesting that for many OGs the presence of the *wild-type* allele is detrimental for the tumor fitness. Additionally, we find additional positive selection pressure on TSG mutations in tumors with CNA gains in the same gene, indicating that dominant-negative mechanisms may be common among TSG point mutations.

### Variant allele frequency distributions shifted by copy-number losses reflect selection

To substantiate that CNA deletions in OGs and also gains on TSGs can increase selection of concomitant mutations, we further considered which allele is affected by the CNA. We hypothesized that mutant allele variant allele frequencies (VAFs) of the driver mutations (here, the assumption is that most mutations in driver gene groups are drivers) will change unidirectionally in samples with either CNA losses or CNA gains, i.e. allelic imbalance will arise (19–21). As a baseline, this is here compared to the random genes set, in which mutations are not selected (i.e. passengers) and so each CNA will presumably affect the mutant allele and the *wild-type* allele with similar frequencies. We corrected VAF estimates for variation in purity between tumor samples (**Methods**), while to account for ploidy effects on VAF, the comparison to a control group of random genes provides a baseline.

In this test, higher VAFs of mutant alleles in sets of cancer genes would be consistent with a stronger positive selection. As a control, we considered the known two-hit TSG mechanism: in tumors with a CNA loss in a TSG, the allele frequencies of nonsynonymous mutations in the TSG in cognate cancer types were significantly higher than those in random genes (Mann-Whitney test test p = 4.34·10^−8^, with the median VAF for TSG = 0.75 and random genes = 0.64) (**Fig. 4A**). In other words, the copy number loss preferentially affected the wild-type allele, resulting in loss-of-heterozygosity (LOH) of the TSG. For the individual TSGs *STK11, NF1, APC*, *VHL, PTEN, CDKN2A, TP53* and others, it was possible to show a significant increase in VAFs in CNA loss states, compared to the set of random genes as a baseline, at FDR ≤ 5% (18 genes significant in total in at least one cancer type at FDR ≤ 25%, see full list in **Supplementary Fig. S7A**).

Next, to support our finding of selective effects of CNA deletions in OGs, we considered VAFs of nonsynonymous mutations therein (**Supplementary Fig. S7B**). Here, the VAFs for missense mutations upon deletion were substantially higher than those of random genes for genes *NRAS*, *KRAS*, *NFE2L2*, *COL3A1, ERBB2* and *TRRAP* at FDR ≤ 25% (**Supplementary Fig. S7B**) Interestingly however, *PIK3CA*, *FGFR3* and *BCLAF1* showed decreased VAFs of missense mutations (FDR ≤ 5%), suggesting haplo-essentiality of those genes, meaning that CNA deletions are not always associated with positive selection in oncogenes, but the effect may be gene or tissue-specific.

To reduce signals of negative selection in OGs confounding this analysis, we analyzed VAFs of missense mutations separately for hotspots and non-hotspot sites. Supporting an increased selection in OGs upon deletion we observed an increased VAFs of oncogenes compared to those of random genes (FDR=25%, p=0.19). This was evident both in VAFs of mutations in strict hotspots, and in non-hotspot mutations in OGs (**Supplementary Fig. S9**, non-significant FDRs). In samples with gene CNA loss, median VAF for OGs in cognate cancer types was 0.59 and 0.63, for non-hotspot and hotspot mutations, respectively. This exceeds the values observed in samples with the known two-hit mechanism linking OGs and CNA gene gain, where the median of the VAF distribution for OGs in cognate cancer types was 0.56 and 0.33 for non-hotspot and hotspot mutations, respectively. Additionally, the VAF of non-hotspot mutations in cognate OGs upon CNA deletions were lower than mutations in strict hotspots (however non-significant difference, **Supplementary Fig. S9**), suggesting an increased effect in hotspots. Considered jointly, this supports that when an OG bears a mutation and a deletion occurs, the wild-type allele tends to be the one deleted rather than the mutant OG allele, signaling selection upon deletions in mutated OGs.

### Copy-number gain associated conditional selection on TSGs and OGs confirmed in VAF analysis

Next, we turn to examine driver mutation VAFs in tumor samples with a CNA gain in OGs and TSGs. In accordance with reports of two-hit events involving gains on OGs (20, 23), the frequencies of missense mutations in OGs showed higher VAFs compared to random genes (Mann-Whitney test test p = 7·10^−19^, with the median VAF for the OGs = 0.65 and random genes = 0.50), suggesting that the mutant allele is the one that gets duplicated more often (**Fig. 4A**). For example, oncogenes *NRAS*, *KRAS*, BRAF, *FGFR3*, *PIK3CA,* and *NFE2L2* individually had significantly higher VAF in all nonsynonymous mutations, when compared to a random genes baseline at FDR ≤ 5%; total number of significant oncogenes with an increased VAFs in samples with CNA gain (in at least one cancer type) reached 20 at FDR ≤ 25% (see **Supplementary Fig. S8B**).

On the contrary, nonsense mutations in OGs in cognate cancer types (with the median VAF for the group of random genes = 0.53 and OGs = 0.4, insignificant difference, Mann-Whitney test test p = 0.47) had lower allele frequencies upon CNA gain, compared to random genes. This indicates there may be a negative selection against the increasing dosage of truncated OG proteins, suggesting that some truncated OGs may act in a dominant-negative manner.

Next, we consider TSG, for which our analyses of conditional selection (see above) suggested that there is, interestingly, overall increased positive selection on point mutations upon gains, thus extending the two-hit TSG paradigm. This is indeed supported also in the VAF analysis: potentially inactivating mutations (both missense and nonsense) had higher VAFs in tumor suppressor genes upon copy number gain, compared to a random genes baseline (**Fig. 4A**). This implies that, when CNA gains occur in TSG, they preferentially amplify the mutant TSG alleles, supporting the findings above about increased selection upon point mutations in TSGs bearing CNA gains. Alternatively or additionally, there may be a negative selection against amplification of wild-type alleles in TSGs bearing mutations. Both of these scenarios are compatible with dominant-negative activity of many mutations in TSGs. Further, our data suggests a model where the dominant-negative mutations in these TSG undergoing CNA gains are only partially dominant, and thus increasing their proportion versus the *wild-type* allele benefits fitness of cancer cells. Some examples of genes associated with CNA gains in this test include *PBRM1*, *STK11*, *CDKN1A*, *TP53*, *CDKN2A*, *NSD1*, *PPP2R1A*, *FBXW7*, *NF1*, *ARHGAP35*, *KEAP1*, *PTPRB*, and *APC* genes, which had significantly higher values of mutant VAFs at FDR≤5% (**Supplementary Fig. S8A**), suggesting an (incompletely) dominant-negative GoF character of at least some mutations in those tumor suppressor genes.

### Replicating associations between selection on point mutations and CNA events in an independent data set

To further verify the positive selection change upon deletion of a gene or copy number gain in TSGs and OGs, we estimated CNA-conditional selection in an independent dataset with 89,243 tumor samples from the Genomics Evidence Neoplasia Information Exchange (GENIE) consortium (50, 51), consisting of panel sequencing data, where we examined 170 genes therein. Here, to estimate a baseline mutation rate neighboring genes were not available, however our MutMatch method used the low-impact nonsynonymous mutations within cancer genes (defined as with CADD<20 (52); **Supplementary Fig. S10**). Additionally, since the sequenced genes are all driver genes, a “random genes” control set is not available but cancer genes in non-cognate cancer types were used as an approximation.

The selection change upon both gain and upon deletion was positive for both the TSGs and OGs, substantiating the overall findings in the discovery cohort. As before, the effect size we measured was the difference (between the diploid state and the CNA state) in (log_2_ fold-enrichment of mutations in the selected region (here, high-CADD) versus the putatively non-selected regions (here, low-CADD gene coding regions). Then, these effect sizes were compared. The difference of distribution medians between the cognate and (as a control) the non-cognate drivers for TSGs was +0.14 and +0.21 for selection change upon CNA deletion and CNA gain, respectively. This replicates the finding from the discovery data set suggesting commonly increased selection upon copy-number gains in TSG.

Next, this same difference of medians for cognate OGs was +0.17 and +0.27 for selection change upon deletion and gain, respectively. This difference observed in independent validation data (note it was not statistically significant) provides supporting evidence of positive selection of the CNA losses in OGs. It substantiates that deletions in OGs commonly act to potentiate selection on oncogenic mutations therein; the effect sizes here were comparable to those resulting from CNA gains, a known two-hit mechanism affecting OGs.

### Main trends in genomic signatures of selection on somatic mutations and CNA in driver genes

Our analysis of ∼17,000 cancer genomes/exomes estimated selection on different types of point mutations (nonsense, nonsynonymous in hotspots, and outside of hotspots) in different copy number states, for every cancer gene and every cancer type. This represents a rich dataset for deriving a systematic classification of cancer genes, by their mechanism of activation or inactivation via combining point mutations and CNA, and by their spectrum of driver potential across many tissues. Previous classifications tended to focus on classifying TSG from OG, and two-hit from one-hit genes (11, 23, 53), while here we consider a multi-dimensional classification for each gene simultaneously considering GoF versus LoF mutations, CNA states, tissue specificity, and interactions of these factors.

To outline general trends in this data on driver gene selection, we performed a principal component (PC) analysis on the estimates of selection (as above, the effect sizes are log_2_ fold-enrichments of mutations in a cancer gene, normalized by the neutral rates estimates) in various mutation types and CNA states and tissues. The aim was to measure covariation between extent of selection on different types of genetic alteration across cancer gene-cancer type combinations, thus inferring “driver mechanism signatures”. A visualization of the first four PCs (**Supplementary Fig. S11**) suggests that cancer genes do not appear to form distinct clusters by the mechanism of (in)activation via mutations and CNA (**Fig. 5A-B**). Instead, we observe a continuous spectrum of cancer-driving potential effectuated via multiple mechanisms (as summarized by each PCs), which act in different relative frequencies on different genes and tissues (**Fig. 4A-C**).

**Figure 5.**
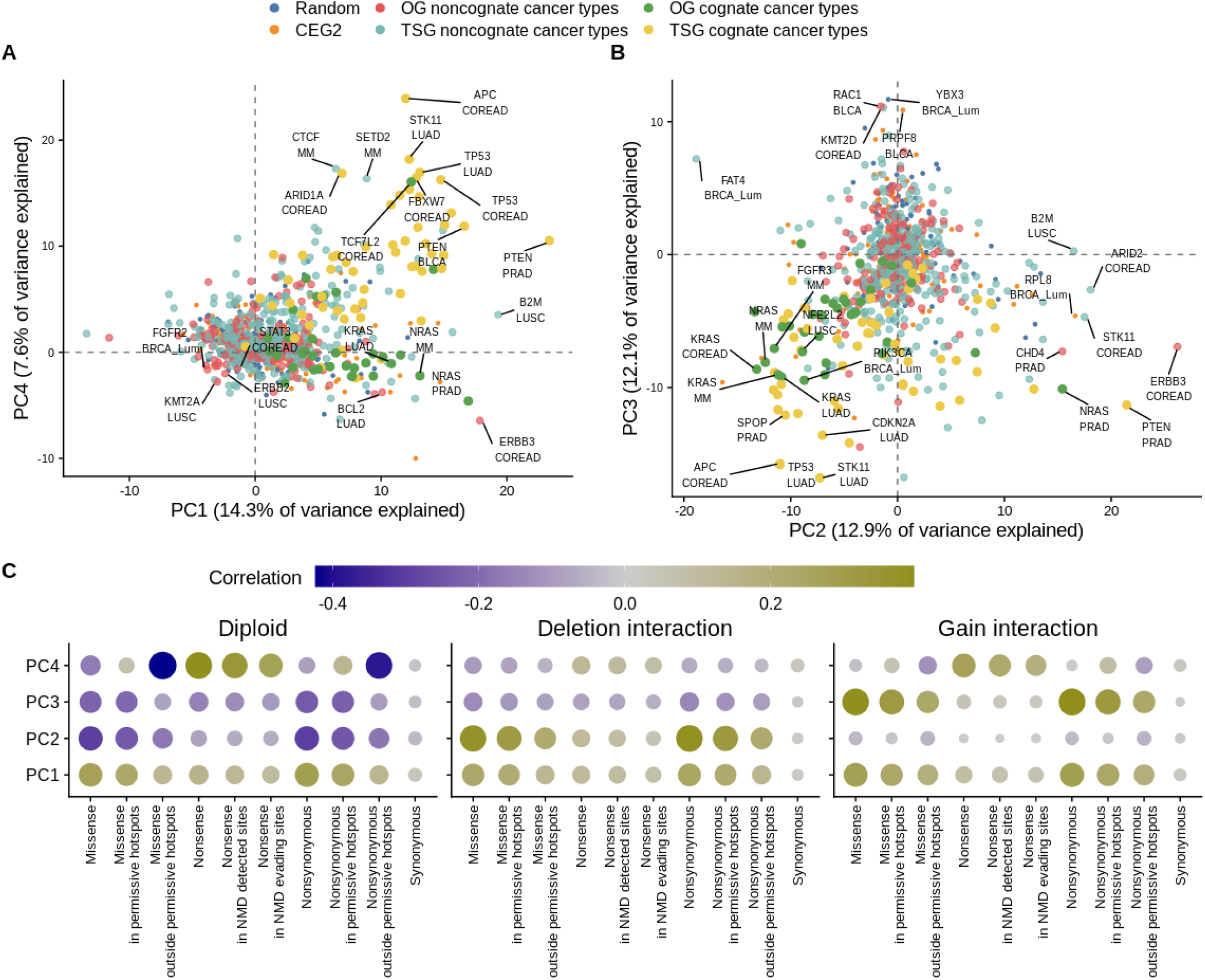
PCA analysis of selection effects across cancer types and mutation classes. Principal components 1 to 4 are shown for 150 most frequently mutated cancer genes and for a subset of random and essential genes. **A-B**. Gene groups in the space defined by the four first PCs. Genes with the highest absolute scores are labeled. **C.** Correlations (loadings) of the principal components with the selection effects for different mutation types.

The most prominent trend in variation between genes reflects simply the overall intensity of positive selection across all copy number states and mutation types (PC1, 14.3 % variance explained): some genes are stronger drivers and others are weaker drivers, which is quantified by PC1 (**Supplementary Fig. S11A-B)**. The following two PCs are informative with regards to interaction of selection on mutations with the CNA state –– PC2 for conditional selection associated with gene loss, and PC3 with gene gain –– and jointly can explain 24.9 % of systematic variance in the selection effects across cancer genes (**Supplementary Fig. S11A-B**). It is notable that these two PCs describing CNA-specific selection effects explain considerably more variance than the general-selection PC1, suggesting that cancer driver genes vary more in their CNA state-specific selection than in the overall selection measured across all CNA states. In other words, changes in driver potential across CNA states are substantial for many cancer genes. Finally, the following signature PC4 accounted for the prevalence of selection of nonsense mutations versus other mutation types, thus separating TSG from OG. This PC4 captures this property commonly used to classify cancer genes — presence of obvious LoF mutations — and explains 7.4 % of systematic variance, which is considerably less than the CNA-interaction PC2 and PC3. Interestingly, PC4 also positively correlated with copy number gains and, to a smaller extent, with gene loss, in agreement with the shown selection of gains and deletions in TSGs. Overall, this PC analysis suggests that genetic alterations causing mutant allele imbalance in driver genes (as per PC2 and PC3, and to some extent PC4) can be very informative features for categorizing cancer genes. Therefore we proceed to generate gene categories based on this data set.

### Deriving a data-driven, comprehensive, mechanistic classification of cancer genes

We next performed another PC analysis on the regularized estimates of selection effects (**Methods**); this results in that the more noisy estimates of selection are brought towards 0, while the more confident ones remain at higher absolute values. Next, we applied hierarchical clustering to the cancer genes described by the resulting 9 factors (PCs to which rotation was applied, see **Methods**). We focused on the 25 cancer types with the largest number of mutations available, and additionally only focused on those gene-tissue pairs where selection effect sizes were non-zero (after regularization, as above).

The driver gene-tissue pairs were classified into 7 clusters by mechanisms of genetic alterations (**Fig. 6A**), the clusters can be further organized into several overarching groups.

**Figure 6.**
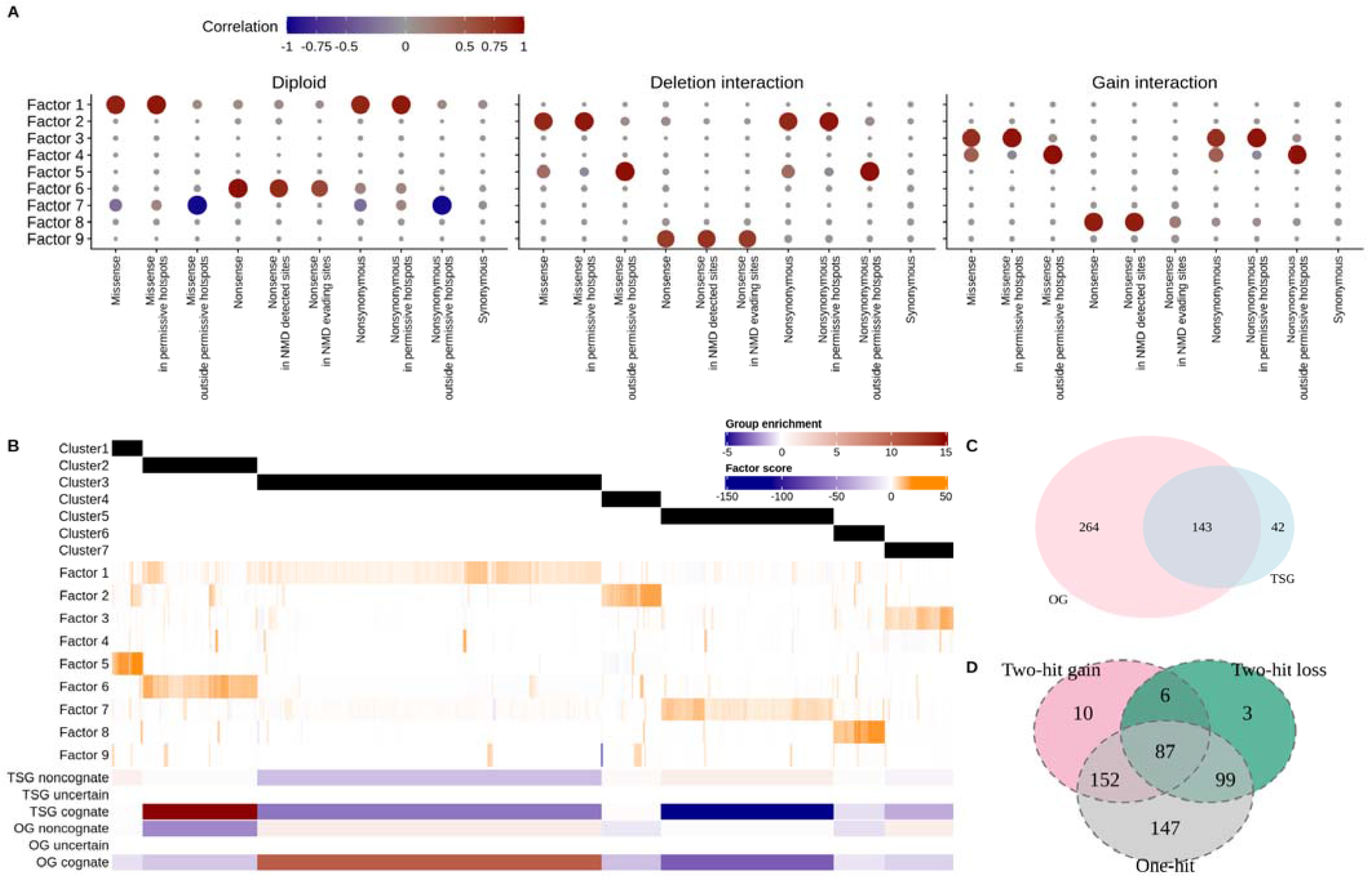
Deriving a data-driven, comprehensive, mechanistic classification of cancer genes. **A**. Correlation between factors (rotated PCs) and original selection effects across mutation types and copy number states. **B.** Clustering of cancer genes across 25 largest cancer types (excluding MSI and ultramutated). Enrichments of gene-tumor group types are shown with signed logarithms of p-values derived from chi-squared residuals, such that deep blue color corresponds to the group depletion, and deep red color corresponds to the group enrichment. **C.** One-hit TSGs and one-hit OGs are not exclusive to their respective clusters, indicating that many genes switch roles in different cancer types. **D.** Cancer genes can switch between one-hit and two-hit depending on the cancer type.

The first group consists of clusters of two-hit genes, with emphasis on CNA losses. This would encompass Cluster 1 (3.7% gene-tissue pairs) and Cluster 4 (7.1% gene-tissue pairs) which are enriched with noncognate genes, thus, presumably weakly selected in the copy number-neutral state (**Fig 6B**).

These two clusters contain two-hit genes with selection acting mostly on missense mutations (non-hotspot mutations or hotspot mutations, depending on the cluster); there is also a two-hit selection on nonsense mutations in Cluster 4.

The second group also consists of two-hit genes, but here with emphasis on CNA gains. The Clu ters 6 and 7 consist of two-hit gain genes, with prevalent selection on nonsense mutations (6.1% gene tissue pairs, with TSGs and also some OGs), or hotspot missense mutations (8.2% gene-tissue pairs, higher enrichment of OGs), in Cluster 6 and Cluster 7 respectively (**Fig 6B**). This Cluster 6 underscores our finding of selection of CNA gains in tumor suppressor genes.

The third group consists of genes that often act via the one-hit mechanisms (as per Factor 1 and Factor 6, **Fig 6A**). These are the Cluster 2 (13.6% gene-tissue pairs) representing one-hit cognate TSGs with prevalent selection on nonsense mutations and missense mutations, and in Cluster 3 (40.9% gene-tissue pairs) representing genes with positive selection on missense hotspot mutations, with enrichment of cognate OGs. The latter cluster of OGs also displays subtle negative selection on nonsense mutations and missense non-hotspot mutations consistent with our analysis above (**Supplementary Fig. S12**). Importantly it is not excluded that some of these genes may additionally act via two-hit mechanisms (with high scores in Factors 3, 4 and 8, see Fig. 6B) however they are classified in this group because of the one-hit effect being prominent.

The fourth group would contain the large, one-hit Cluster 5 (20.5% gene-tissue pairs), which is a mixed category consisting of weakly positively selected genes, both TSGs and OGs. However unlike Cluster 1 and Cluster 4 there is not a detectable two-hit selection on these genes, thus selection on both the copy-number neutral and the CNA gained/lost states would be similarly subtle. We observed increased scores of Factor 7 in this cluster (**Fig. 6A**, **Supplementary Fig. S12**), which is correlated with negative selection of non-hotspot missense mutations, indicating that this cluster contains genes that are essential for tumors via a hypothetical “noncanonical addiction” mechanism. Some genes may switch between the OG clusters and the TSG clusters, depending on the cancer type (**Fig. 6C**).

Another way of classifying these 7 driver gene clusters is by one-hit genes (Clusters 2, 3 and 5) versus two-hit genes (Clusters 1, 4, 6 and 7). Considering the genes within these clusters suggests the majority of the genes may switch between one– and two-hit mechanisms between different cancer types, consistent with a recent study (23), while the minority of the cancer genes (29%) being predominantly one-hit genes, or in contrast 3.8% being predominantly two-hit genes (**Fig. 6D**).

In summary, our analysis delineated 7 major cancer gene categories, providing a data-driven, unbiased classification of driver genes. These categories exhibited different selection pressures on different mutation types and copy number states, and different spectrum of affected cancer types.

## DISCUSSION

The prevalence and the strength of the selection acting to purge deleterious somatic mutations in cancer cells have been unclear. It was clear, though, there is an interaction thereof with the copy-number state of genes (1, 28, 31). Thus we considered both variables jointly to interrogate signatures of negative selection acting on essential genes and also on cancer genes. We suggest that point mutations in cancer genomes can be both positively and negatively selected in the same genes, as we find is commonly the case with oncogenes, in which we estimate ∼20% of point mutations may be purged. Previous studies suggested negative selection against specifically nonsense mutations in OGs (27, 29) (possibly, a genomic signature of oncogene addiction (54)), consistent with the fact that the function of the OG is vital for the survival and proliferation of cancer cells. Quantifying this negative selection in oncogenes may have implications to identifying cancer vulnerabilities, particularly in light of our data suggesting finding that OGs may be negatively selected, and so constitute therapeutic targets even in some cancer types where they are not commonly bearing driver mutations. Based on genomic signatures, we have tentatively named this prediction “non-canonical oncogene addiction”, however experimental validation of this awaits. A limitation of our statistical genomic analysis is that it is not powered to detect negative selection on individual OGs, thereby identifying particular candidate vulnerabilities; larger datasets would be expected to remedy this.

With respect to core essential genes (34), our methodology suggests that mutations in them were depleted compared to a mutation rate baseline (here derived from select neighboring genes and by further matching by trinucleotide composition). Consistent with previous work (1, 31), the effect size of this negative selection is relatively small — we estimate it removes ∼10% mutations from a set of known essential genes — therefore it can be detected only in very large datasets, such as our pan-cancer analysis with ∼18 000 somatic genomes. Moreover our mutation rate baseline, which controlled for effects of concurrent CNA on apparent mutation rates, may have facilitated detecting the more subtle signal. While identifying individual genes under negative selection is of high interest for identifying novel cancer dependencies, substantially larger sample sizes would be needed to gain statistical power to make this task feasible.

Our analyses suggest that the net distribution of mutations in OGs results from simultaneous activity of positive selection that increases the frequency of cancer-driving mutations, and purifying selection that removes the deleterious mutations. The canceling out of signals from positively and negatively selected mutations can lead to underestimation of the positive selection strength. We suggest a very simple method to circumvent this and increase power to detect less frequently-mutated “long-tail” driver OGs in certain cancer types, by focussing the selection test on the regions that are a priori known to be likely under positive selection. This is related to existing approaches that test for mutation clustering in cancer (44, 55) and is limited to known driver genes, however it does identify additional cancer types driven by a gene, better describing the tissue spectrum for each gene.

We comprehensively analyzed the relationships between selection on point mutations and the CNA events at the same locus. One interesting observation concerns two-hit mechanisms, which are well established for TSGs and deletions, but were also reported in OGs in gains/amplifications (20, 23, 56). We suggest that drawing on this may be used to identify additional selected driver genes relevant to a certain cancer type, by testing for additional selection in tumors bearing CNA, potentially boosting signal-to-noise in the analysis; we report various plausible examples of genes that would be missed in the general selection test but would be identified by focussing on CNA-altered states.

Further, our observations clearly implicate not only gains, but also CNA losses as associated with increased positive selection on mutations in OGs. This is anticipated by an analysis of allelic imbalances at the DNA level in oncogenes, which associate not only with CNA gains but also losses (24). Based on this, we suggest that many mutations in OGs may not be fully dominant, but necessitate a two-hit mechanism for increased fitness benefits to tumor cells. Classically, this would be via a CNA gain however we propose a deletion may be another path towards the same goal. Thus, our data provides genomic evidence that the wild-type allele of OG can have a negative effect on tumor fitness, as previously shown for the case of RAS genes (24, 57–59) in particular, and so positive selection often acts on reducing dosage of the *wild-type* OG allele of some OGs.

Tumor suppressive effects of a TSGs can be lowered by deletion of the wild-type allele, as per classical two-hit mechanism (18, 19). However we find genomic evidence that a fitness benefit is also commonly obtained in TSGs by gaining an additional mutant gene copy via CNA. This suggests that many of the TSG mutations act in a dominant-negative manner, and also that they are incompletely dominant. Thus two-hit mechanisms are prevalent in TSG, and, interestingly, are often effectuated via copy number gains, where the mutant allele tends to be the one undergoing gains.

However, our analysis also suggests there is some nuance to TSG inactivation mechanisms: while two-hit is common (incl. with CNA gains), only a few TSGs in cognate cancer types have an exclusively two-hit mechanism of inactivation i.e. it is rare that TSGs show no or weak selection of mutations in the diploid state. In other words, TSGs are often both one-hit (haploinsufficient or dominant-negative) and also two-hit genes and that these two mechanisms are not mutually exclusive. Some fitness would be gained by the first hit on the TSG and additional fitness would be gained by the second hit, suggesting a gradual inactivation model.

Our global analysis of selection effect sizes in different copy number states in cancer genomes suggest that cancer genes do not form well-separated clusters of two-hit genes versus one-hit genes or selected genes versus not selected genes. Rather, cancer genes span the spectrum between one-hit and two-hit mechanisms of (in)activation, meaning that both mechanisms are operative in the same gene with varying frequencies across tumor types.

## MATERIALS AND METHODS

### Modelling mutation rates with the MutMatch approach to obtain selection effect estimates

To describe the variability in mutation counts in a genomic locus, we model raw mutation counts Y using the following generalized linear model, regularizing by using a weakly informative prior distribution of

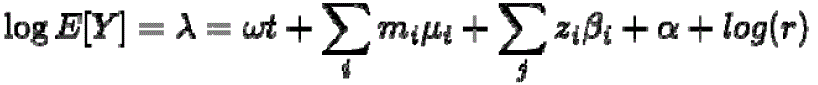

regression coefficients (60):

(1)

where **t** is a target variable used to distinguish mutations accumulated in a genomic area that is currently being tested for a selection signal (central gene) and the control group (neighboring genes). The coefficient ω (or selection effect) reflects the log-fold change of mutation rates in the tested genomic area (where **t** = 1) compared to the region used to model a baseline mutation rate (where **t** = 0). Positive ω estimate indicates enrichment of mutations and positive selection, while negative ω estimate corresponds to the mutation depletion in the area of interest and negative selection.

To control for the different activities of mutational processes on different oligonucleotides (1, 4, 61), the model stratifies mutational counts according to the 96 mutation spectra in trinucleotide context (MS96) using the mutation type variables m and corresponding effects μ. Other types of optional variables can be included to control for inter-cancer type differences in selection (in a pan-cancer analysis), confounders or batch effects (such as the source of the data). We denote these variables as **z** and their corresponding effects as β. The base mutation rate α is included as the intercept of the model. Finally, to adjust the mutation counts to the maximal number of “opportunities” for mutations, we include the number of nucleotides-at-risk **r** as an exposure term. This allows us to model rates using mutation counts i.e..it accounts for different DNA length in genomic loci and in trinucleotide contexts.

When testing for conditional selection signals in genes we use an extended version of the regression that includes a condition variable **c** encoding the state of the genomic region with respect to the condition (here, copy number state of a gene) and the interaction term of the target variable **t** with the condition

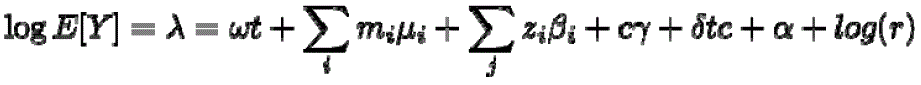

variable **c**.

(2)

To exclude the effect of mutation rate heterogeneity at the gene-level (i.e. within-neighborhood variation) that can confound the framework, we exclude genes in the neighborhood that have a different mutation profile than the central gene, quantified by the outlier score S_outlier_

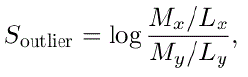

where M is the number of mutations observed in a gene (central gene or a gene from the neighborhood), and L is the gene length. Mutation counts of individual genes for this outlier analysis were obtained from intergenic and intronic regions within the +-20kb of the center of each gene, using WGS data processed in Salvadores *et al.* (62), where the control for trinucleotide content was applied with minor adaptations (matching tolerance threshold ≤0.035 with 10 000 iterations). 11 genes without mutations (after a matching procedure) were excluded to avoid infinite values of S_outlier_. In the end, an S outlier value was calculated for 18 214 transcripts. For each gene, neighborhood genes with |S_outlier_| > 0.2 were excluded, thus allowing less than 22% variance between the central gene and genes in the neighborhood.

For each gene, only nucleotides of coding exonic sequences of the most expressed transcript (63) (with adjacent 5 nucleotides upstream and downstream of each exon to account for splice sites) that are mappable according to the CRG75 Alignability track (64) were considered. Additionally, we removed positions that were unstable when converting between GRCh37 and GRCh38 (conversion-unstable positions).

To remove biases in selection estimates caused by the low mutation counts in a region, we generated a null distribution of selection estimates for ω and δ coefficients using a randomization procedure. The mutations observed in each cohort are randomly shuffled between the tested and control genomic loci, with a chance of acquiring mutations proportional to the number of sites in the loci. Then, the same selection model was applied to the randomized data (using 50 repetitions) to estimate the null distribution of selection effects. We subtract the median of this null distribution for a coefficient from the coefficients estimates obtained from the actual data for the later analysis.

To obtain selection estimates across all copy number states, we considered all samples together without stratifying into different copy number states as in (1), and not controlling for any additional factors z. To obtain selection estimates for the conditional selection cohort, we performed two separate analyses as in (2) with the condition variable **c** varying between analyses. One regression included diploid samples versus samples with CNA loss, another regression with diploid samples against samples with CNA gain. Estimation of conditional selection in the GENIE validation cohort was performed similarly, with an additional variable denoting the cohort (DFCI or MSK) to control for possible batch effects between cohorts. In case of pan-cancer selection estimation, cancer type was additionally included in the model.

Finally, we only focused on genes that had at least 2 (4) mutations across all samples where a gene was in a diploid state and at least 2 (4) mutations in samples where a gene copy was deleted or gained, separately for each cancer type for the discovery and validation datasets, respectively. For the pan-cancer analysis we required at least 10 mutations to occur in the gene of interest across all cancer types used, and for selection estimates across copy number states we required at least 8 mutations to be present in each regression model.

Multiple testing FDR correction was performed using the Holm method separately for each cancer type and group of genes (CEG2, TSG in cognate cancer types, OG cognate cancer types, TSG in noncognate cancer types, OG noncognate cancer types and random genes).

### Mutation and copy number data collection and processing

We collected mutation and copy number data for two aggregated datasets in this study: a discovery data set and a validation data set. The discovery data comprised WES and WGS datasets from WES somatic Single Nucleotide Variants (SNVs) from the The MultiCenter Mutation Calling in Multiple Cancers (MC3) Project (65), WGS somatic SNVs from the TCGA consortium (66), WGS somatic SNVs from Hartwig Medical Foundation (HMF) project (67), WGS somatic SNVs from Pan-Cancer Analysis Of Whole Genomes (PCAWG) dataset (68), WGS somatic SNVs from Personal Oncogenomics project (POG570) program (69), WES somatic SNVs from The Clinical Proteomic Tumor Analysis Consortium (CPTAC)-3 program (70, 71), WGS somatic SNVs from MMRF-COMPASS study (72). The mutations that were called against the human version assembly GRCh38 were converted to the hg19 reference genome using LiftOver (73). Variant (nonsense, missense, synonymous) were annotated using ANNOVAR software (74). Ultramutated samples, and samples with a high fraction of indels were separated from the rest of the samples for UCEC, CESC, COREAD, ESAD, ESCA, STAD, UCS cancers into separate subtypes denoted as “POLE” or “MSI” (Table 9.1). Cancer types between different datasets were matched to increase the sample size for each of them.

Altogether, we collected the genomes of 23 000 tumor samples with mutations from 117 cancer types; the number of tumor samples per each tumor type was ranging from 1 to over 1000 for PRAD (prostate adenocarcinoma), BRCA-Lum (a luminal subtype of breast cancer), COREAD (colorectal adenocarcinoma), MM (multiple myeloma), kidney, PAAD (pancreatic adenocarcinoma), and SKCM (skin cutaneous melanoma). The median number of tumor samples per cancer type was 57. The average number of SNV mutations in each cancer type was 28 805, and the median number of mutations was 4 136. The cancer types with the biggest number of mutations are listed in **Supplementary Table 1**.

For the validation dataset, we downloaded mutation calls for 90 713 tumor samples across 75 cancer types from MSK-IMPACT and the Dana Farber Cancer Institute (DFCI) Oncopanel of the American Association for Cancer Research Project Genomics Evidence Neoplasia Information Exchange (GENIE) (Release 11.1; syn32309524) (50, 51). In these studies, only a limited number of cancer genes were sequenced. We determined the list of cancer genes that were targeted in both of these cohorts.

We collected Copy Number Alteration (CNA) data estimated with GISTIC2, ASCAT or purple programs for the majority of the samples listed above, covering 17 644 samples of the discovery cohort and 89 243 samples of the validation cohort. The estimates of the gene-level copy number status were binarized according to the recommended sample-specific thresholds for the GISTIC2 copy number levels (12). For the discovery cohort, only low-level gains or hemizygous deletions were considered for estimation of conditional selection upon CNA alteration event, while high-level amplifications and homozygous deletions were not considered. For the validation cohort, we have aggregated the estimates for the copy number status of the gene such that any sample with the number of gene copies greater than 2 was considered to be in a gain state, and any sample with the number of gene copies less than 2 was considered to be in a deleted state.

### Annotation of cognate cancer types

To determine which cancer types were cognate (i.e. where cancer genes are positively selected), we used different sources of data: annotation available in the known databases such as Cancer Gene Census (CGC) or MutPanning list (9, 75), and additionally as the main criterion we used the selection estimates that we derived using MutMatch with the neighboring genes baseline in the discovery dataset of WES/WGS tumor. The latter approach (our annotations based on MutMatch) carries fewer risks of missing cognate cancer types due to the different labels between annotation sources and our data, or uncertainty in the case when annotations are not detailed to specify which cancer subtype is cognate. Moreover, this helps to make sure that we do not focus on gene-tumor pairs that are listed as cognate, but may have too few mutations in datasets analyzed here due to the low number of samples in these cancer types.

To infer which gene-cancer type pairs are cognate, we have estimated the selection effects in the discovery data set for all the cancer genes in the analysis across all copy number states (in this case, without controlling for the CNA variable). Cancer types where a gene was positively selected (ω > 0 at a threshold of FDR ≤ 25 %) were considered to be cognate cancer types for this gene. To infer noncognate cancer types (where cancer genes are not strongly selected) false-negative hits, we required FDR ≥ 75 %. The rest of the gene-cancer type pairs that did not fall into either of these categories were separated into the “uncertain” group.

### CADD scores

We downloaded a bigWig file that contained pre-computed PHRED-like −10 × log 10 rank total scaled Combined Annotation-Dependent Depletion (CADD) scores for each genomic position (v.1.4) (52). The highest CADD score of any 3 possible substitutions at a site is provided in this file, with higher values indicating a higher level of deleteriousness of the variant. Scaled CADD-scores assign value 10 to the top-10 % of all the CADD scores in the reference genome, value 30 to the top-0.1 % and so on. To separate regions where mutations are likely to have a functional impact versus regions that are evolutionary unconstrained, we used a cut-off of 20. In this way, positions in a gene with top-1 % most deleterious mutations were tested for selection using a background mutation rate model from unconstrained gene regions where CADD score was less than 20.

### Essentiality scores

We used mean cell-essentiality scores across different cell lines (CERES score by CRISPR-Cas9 approach (47)) and essentiality score derived from a population data (selection of Loss-of-Function (LoF) germline point mutations summarized in Loss-of-function Observed over Expected Upper bound Fraction (LOEUF) score (48)) to independently define cell line-essential and population genomics-essential genes.

### Hotspots

We defined “permissive hotspots” based on the codon-specific frequency of mutations in a discovery cohort with the cut-off of 2 mutations per codon, which corresponds to the 91.6 % of recovery of Trevino hotspots and 59 % of recovery of 3D hotspots (76) (0.235 % and 0.007 % of all permissive hotspot sites, respectively). Some 3D clustered hotspots that were not recovered with this cut-off are located in proximity to a hotspot in the discovery cohort. To avoid missing such sites that are closely located to a detected hotspot, but have fewer mutations (due to the limited number of mutations in the discovery dataset), we lengthened our hotspots to include them. If another mutation was found within 3 codons upstream or downstream from the hotspot, the hotspot was extended to include this mutation. The median length of the hotspots defined this way was 12 nucleotides, or 4 amino acids. This recovered 75.2% of 3D clustered hotspots, and 95.4% of Trevino hotspots (40).

### NMD-detected and NMD-evading regions

Genomic regions were split into those where premature termination codons (PTCs) lead degradation of the mRNA in a process of nonsense-mediated mRNA decay (NMD) or those o the where nonsense mutations lead to a translation of a truncated protein sequence. The efficacy of NMD for PTCs in a human model was predicted using the NMDetective algorithm (77). Regions with the NMDetective-A score >0.52 were classified as NMD-detected regions, and the rest were classified as NMD-evading regions.

### VAF analysis

VAF analysis was performed using the TCGA dataset. We calculated allele frequencies for missense and nonsense mutations located in the same genomic regions that were used for the estimation of selection (including removal of unmappable and conversion-unstable positions). VAF estimates were then adjusted to control for a sample purity (Consensus measurement of Purity Estimation from TCGAbiolinks (78):

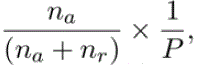

where n a and n r is the number of alternative and reference reads, and P is the sample purity.

### PCA on selection estimates

The de-biased estimates for (i) selection effect in the diploid state ω, (ii) selection effect change upon deletion δ_deletion_, and (iii) selection effect change upon gain δ_gain_, for each gene in each cancer typ were used to perform a global analysis of copy-number dependent selection in the human soma. To reduce the likelihood of selection in the control group of random genes, we excluded from them genes with low LOEUF score (ranked in the most essential 30% of the genes) derived from population variants (48, 79), CEG2 essential genes (34) and cancer genes using MutPanning and CGC gene lists (9, 41, 75). We also excluded genes that were either very short or very long, allowing up to 30% of difference between the median number of the nucleotides in a gene that was used in the regression model. Finally, we removed genes with very high or very low gene expression using the normalized transcript expression levels summarized per gene in 54 tissues based on transcriptomics data from Human Protein Atlas (HPA) and Genotype-Tissue Expression (GTEx) (80).

For the selection in the diploid state ω, we averaged the estimates for the diploid-state selection obtained from two different models: one for gene loss and another for gene gain. In this analysis, we focused on cancer types and the genes for which selection effects were estimated in at least 85% of gene-tumor type pairs i.e. common driver events. Because of the high number of missing values in the case of selection estimates for strict (Trevino) hotspots, we excluded estimates inside or outside of the strict hotspots from the PCA analysis (the permissive hotspot estimates were still included). For the gene-tumor type combination passing these filters, missing values were imputed using *imputePCA* function from the “missMDA” package in R.

PCA was performed on the centered and scaled data. To avoid overplotting, we only show 15 genes from the control group of random genes and 15 genes from the control group of essential genes in the main figure.

### Clustering of cancer genes in different cancer types

We performed centered and scaled PCA on the restrained and corrected coefficients to reduce the impact of outliers. To do so, we first forced the highest absolute estimates to be within 98% of all estimates (extreme estimates of top and bottom 1% were capped at the level of 1st percentile or 99th percentile). Next, we subtracted one standard error from each positive estimate (or added it to the negative ones) to obtain more conservative estimates of the selection for each gene-tumor pair.

Missing values were imputed using *imputePCA* function from the “missMDA’’ package in R. We chose the first 9 PCs based on the scree plot for further classification and performed a rotation of PCs using the varimax method to ease the interpretability of obtained factors. We clustered gene-tumor pairs for 25 largest cancer types excluding hypermutated and MSI samples using partitioning around medoids (*pam* function from package “cluster” in R) based on euclidean distances between samples. We performed two clusterings: first yielded 8 clusters (silhouette width criterion), with the largest one corresponding to the pairs showing no selection (factor scores ∼0). To refine this analysis, we excluded pairs from this cluster and performed another clustering, finally discovering 7 clusters of gene-tumor pairs. Enrichments of particular gene classes for each cluster were estimated using two-sided p-values derived from chi-squared residuals (*chisqtest* function from “zebu” package in R).

## DATA AVAILABILITY

The data and code generated in this study is available from the authors upon request. Previously published data and resources used in this work: vcf files and CNV data for TCGA WGS [https://portal.gdc.cancer.gov/], HMF [https://www.hartwigmedicalfoundation.nl/en/], TCGA WES mutation calls from MC3 [https://gdc.cancer.gov/about-data/publications/mc3-2017], TCGA WES CNA calls [https://portal.gdc.cancer.gov/], PCAWG [https://dcc.icgc.org/pcawg/], POG570 [https://www.bcgsc.ca/downloads/POG570/], GENIE [https://www.synapse.org/], Project Score CRISPR genetic screening data [https://depmap.org/portal/download/], gnomAD [https://gnomad.broadinstitute.org/downloads], CADD scores [https://krishna.gs.washington.edu/download/CADD/bigWig/], NMDetective [https://figshare.com/articles/dataset/NMDetective/7803398], CRG75 Alignability track [https://hgdownload.soe.ucsc.edu/goldenPath/hg19/database/], CUP bed files [https://github.com/cathaloruaidh/genomeBuildConversion/#2-novel-cup-bed-files], Top-rank expressed transcripts of protein-coding genes [https://tregt.ibms.sinica.edu.tw/index.php#tab6].

## ACKNOWLEDGEMENTS AND FUNDING SOURCES

We are grateful to Marina Salvadores for assistance with determining mutation burdens of genes. Work in the F.S. lab is supported by an ERC StG “HYPER-INSIGHT” (757700), Horizon2020 project “DECIDER” (965193), Spanish government project “REPAIRSCAPE”, CaixaResearch project “POTENT-IMMUNO” (HR22-00402), an ICREA professorship to F.S., the SGR funding of the Catalan government, and the Severo Ochoa centers of excellence award of the Spanish government to the hosting institution. E.B. was funded by a AGAUR-FI fellowship of the Catalan government. This publication and the underlying research are partly facilitated by Hartwig Medical Foundation and the Center for Personalized Cancer Treatment (CPCT) which have generated, analyzed and made available data for this research (request number DR-260).

