## Supplementary Figures S1-S12 for "Copy number losses of oncogenes and gains of tumor suppressor genes generate common driver events of human cancer"

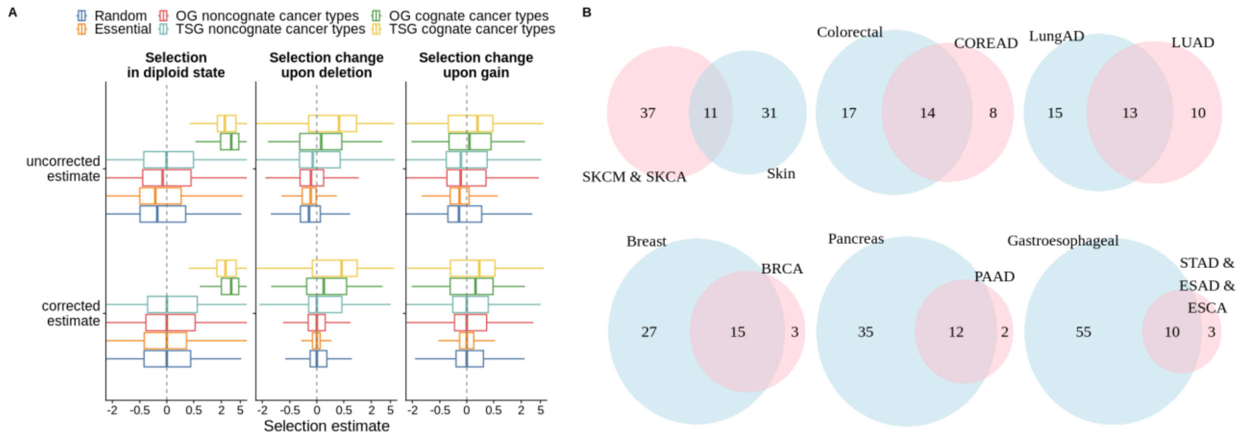


**Supplementary Figure S1. Correction procedure removes a bias from selection estimates and overlap between annotation of driver gene-tumor pairs and known driver genes according to MutPanning. A.** Selection estimates before and after correction. Estimates for selection on nonsynonymous mutations in a diploid state and a conditional selection in a hemizygous state or cells with copy number gain. Uncorrected estimates tend to be more negative than expected, based on the assumption that random genes should be under neutral selection. Bias-corrected estimates for random genes are centered at zero for all copy number states, as expected. **B.** Overlap between selected genes according to the MutPanning method (in blue) and cognate genes defined in this study (in pink) for several cancer types.

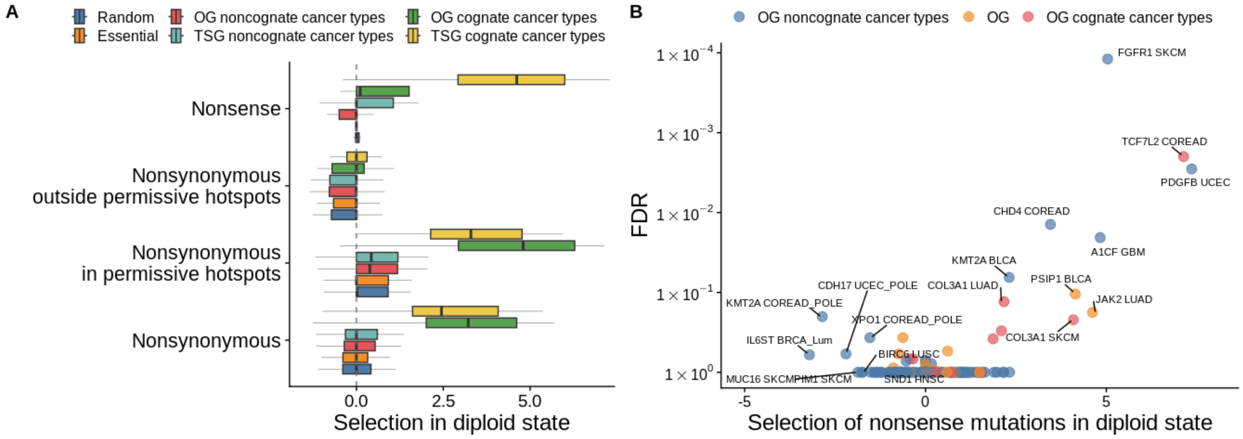


**Supplementary Figure S2. Selection estimates for diploid state show signatures of positive and negative selection in oncogenes. A.** Selection in diploid state. Debiased selection estimates for the neutral copy number status of a gene obtained on the discovery cohort. The number of gene-tumor pairs used to produce each of the box plots is written in the left part of the plot. One data point corresponds to one gene-tumor combination. **B.** Selection of nonsense mutations in the diploid state for oncogenes.

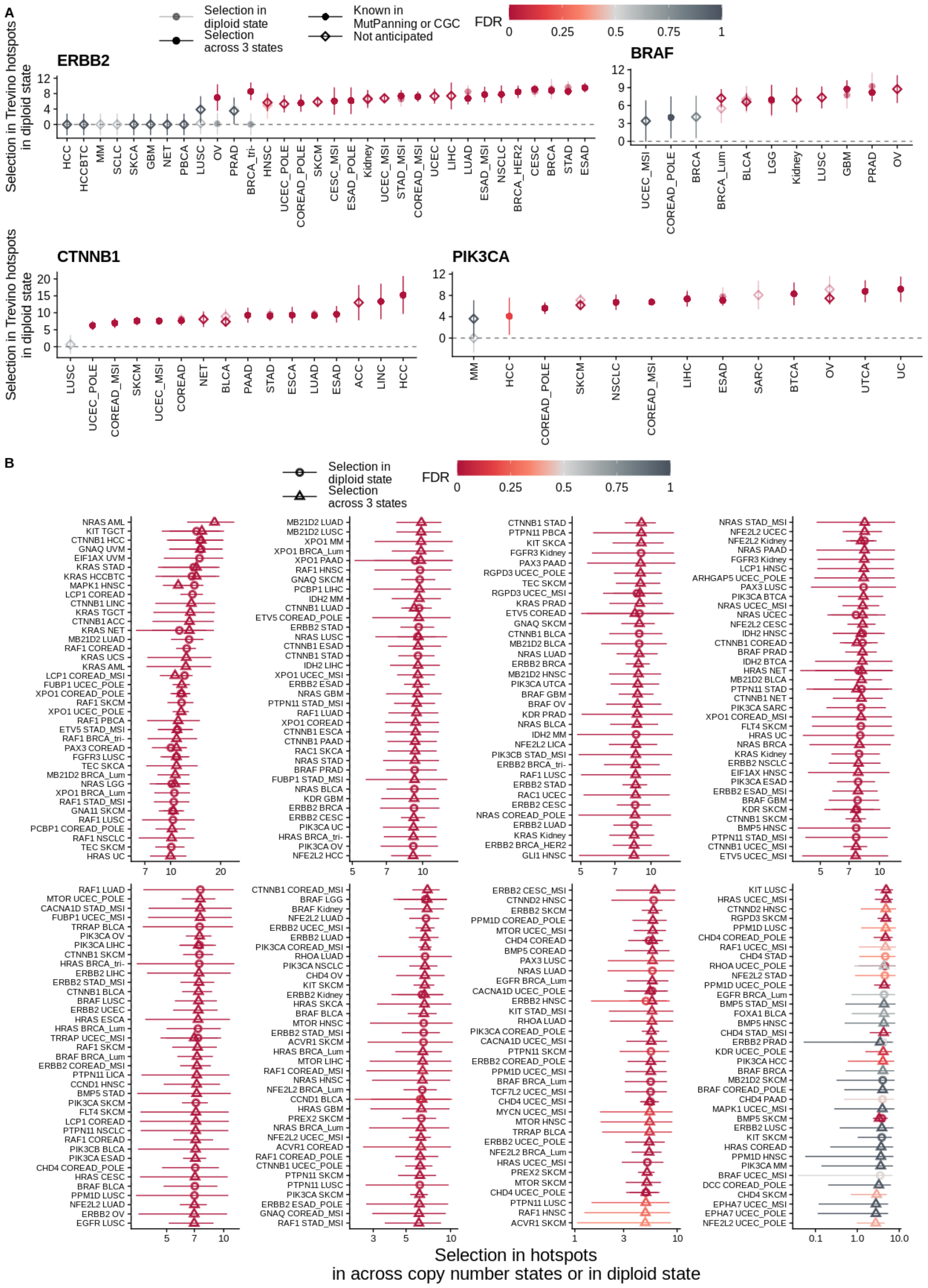


**Supplementary Figure S3. Selection in hotspots helps identify new cognate gene-tumor type combinations. A.** Selection of *ERBB2*, *BRAF*, *CTNNB1* and *PIK3CA* genes in noncognate cancer types. Cancer types where a gene is known to be a driver according to MutPanning or CGC annotations are marked. **B.** Hotspot election in OGs in noncognate cancer types.

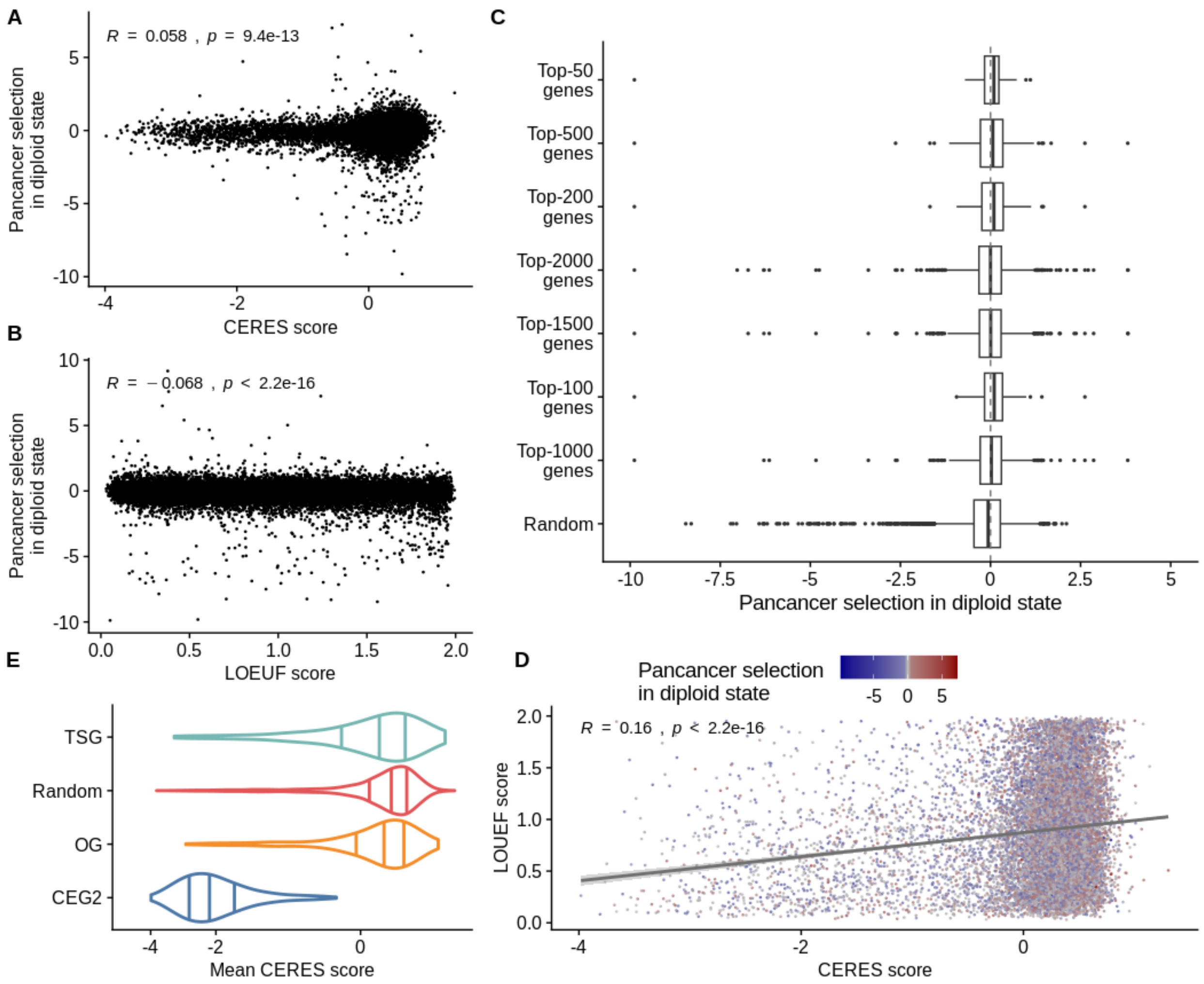


**Supplementary Figure S4. Whole-genome analysis of pan-cancer selection. A-B.** Somatic cell lines derived and population essentiality scores do not correlate with somatic selection estimated in tumors. Lower values of CERES and LOEUF scores correlate with a higher chance of being essential in cell-line knockout experiments or with haploinsufficiency at the population level. We observe very weak correlation between these essentiality scores and estimates of selection derived with the MutMatch method (in the diploid state, pan-cancer analysis) across all genes (Pearson's R=0.06 for CERES score and -0.07 for LOEUF). **C.** The most essential genes according to the LOEUF scores are not more negatively selected than a set of random genes. **D**. LOEUF and CERES scores are weakly correlated (R=0.15). **E**. CERES scores distribution, where lower scores correspond to higher cell essentiality in CRISPR–Cas9 genetic screens.

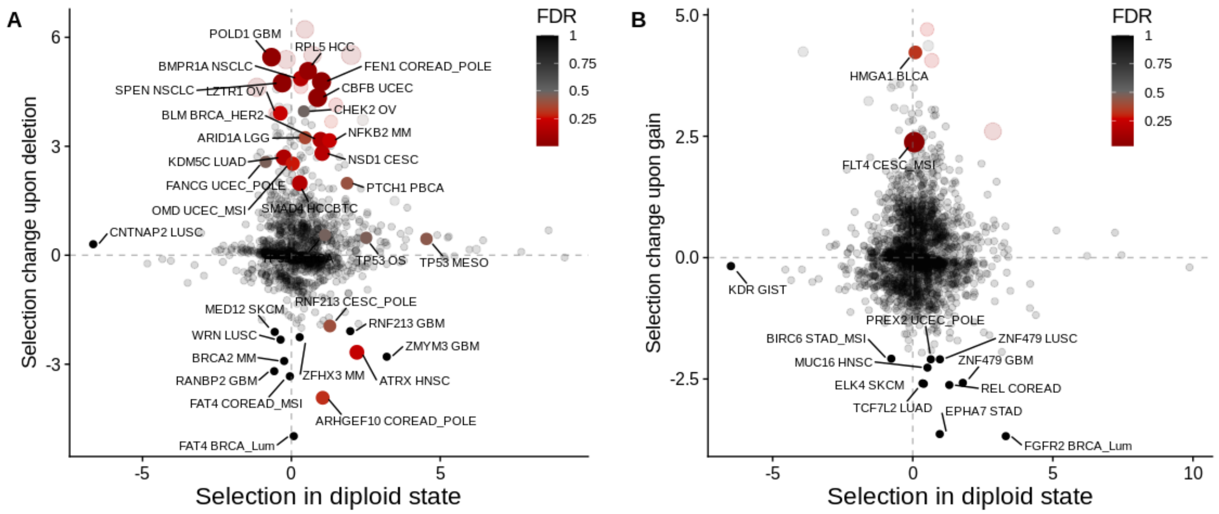


**Supplementary Figure S5. Noncognate gene-tumor combinations selected as two-hit genes.** Selection effects of nonsynonymous mutations in cancer genes in noncognate cancer types: **(A)** TSGs **(B)** OGs. Genes with the strongest selection change in samples with deletion or gain are labelled.

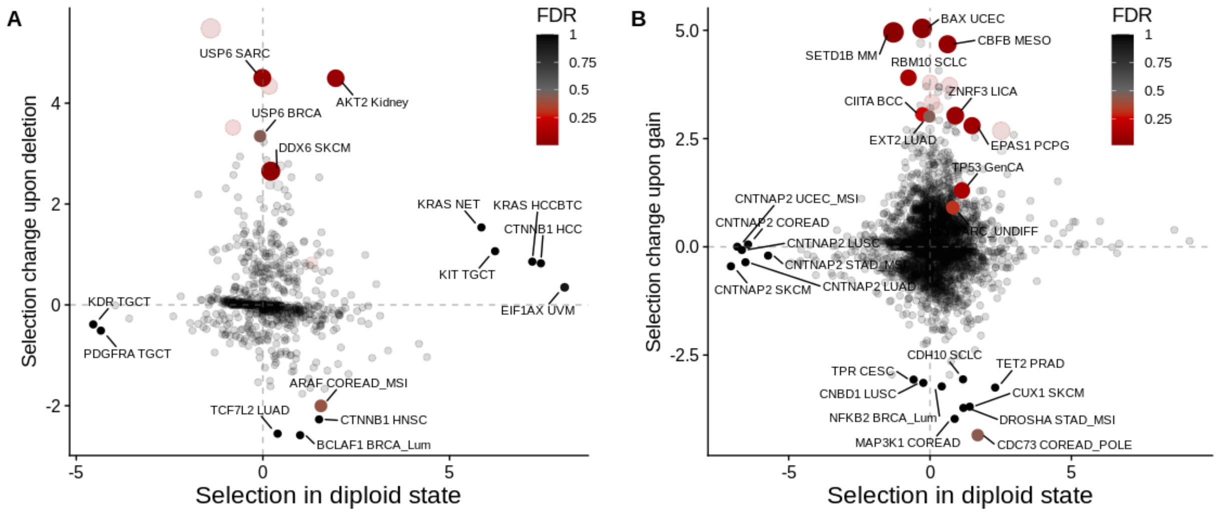


**Supplementary Figure S6. Noncognate gene-tumor combinations selected as two-hit genes.** Selection effects of nonsynonymous mutations in cancer genes in noncognate cancer types: **(A)** OGs **(B)** TSGs. Genes with the strongest selection change in samples with deletion or gain are labelled.

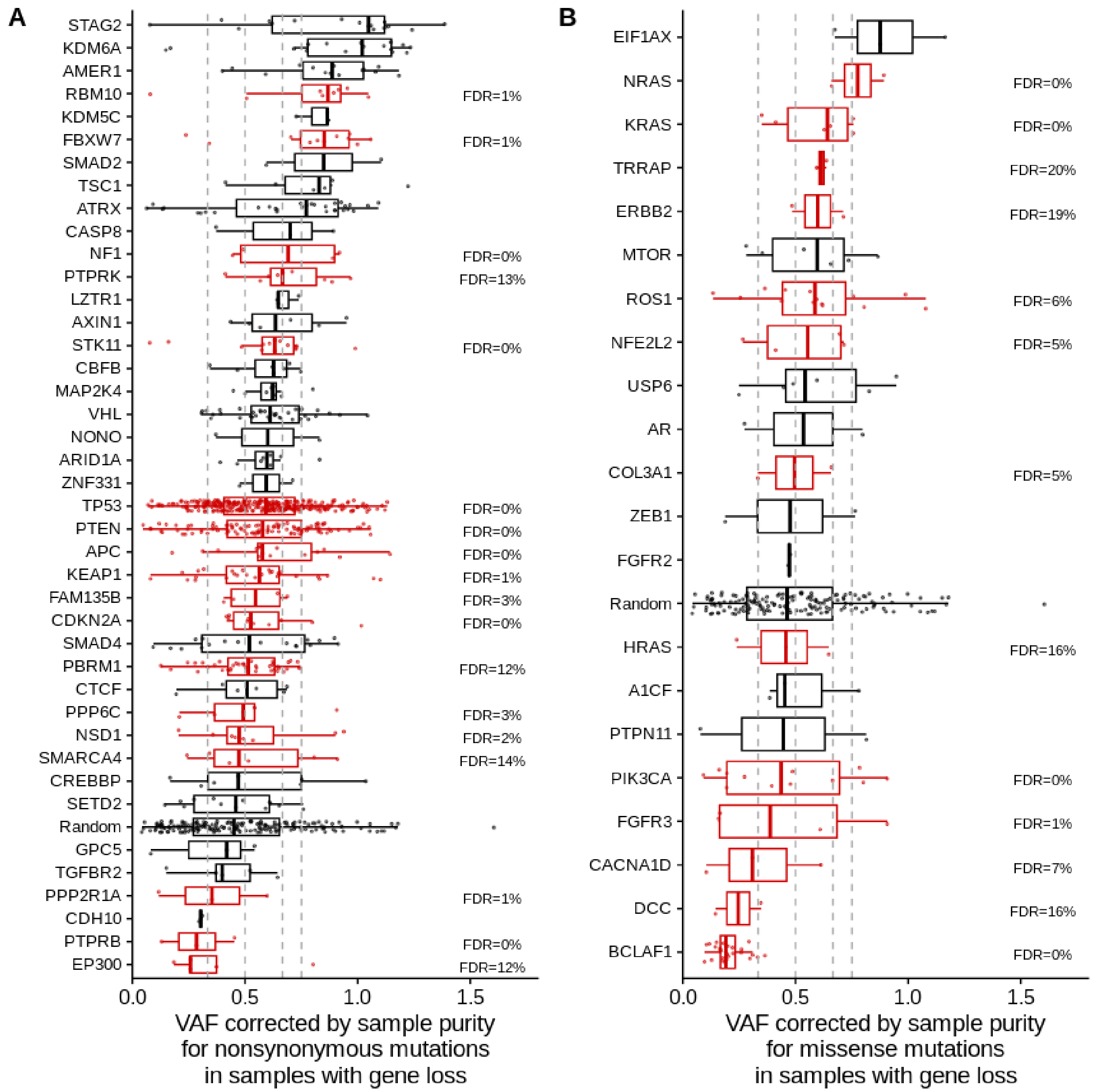


**Supplementary Figure S7. Mutation frequencies in samples with a gene loss. A.** Variant allele frequencies corrected by the sample purity for TSGs across cognate cancer types (one data point corresponds to one nonsynonymous mutation). **B.** Variant allele frequencies corrected by the sample purity for OGs across cognate cancer types.

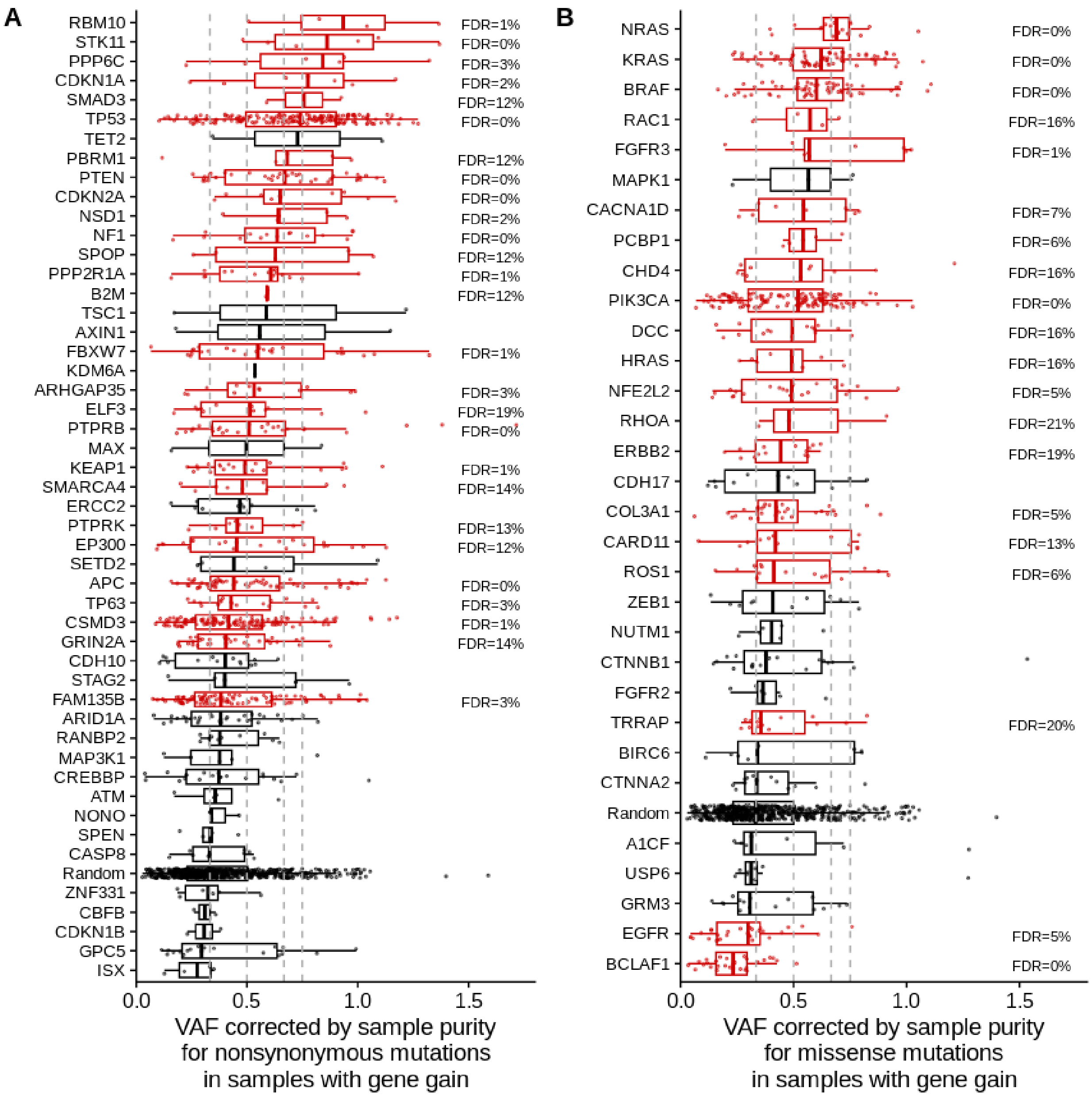


**Supplementary Figure S8**. **Mutation frequencies in samples with a gene copy number gain. A.** Variant allele frequencies corrected by the sample purity for TSGs across cognate cancer types (one data point corresponds to one nonsynonymous mutation). **B.** Variant allele frequencies corrected by the sample purity for OGs across cognate cancer types.

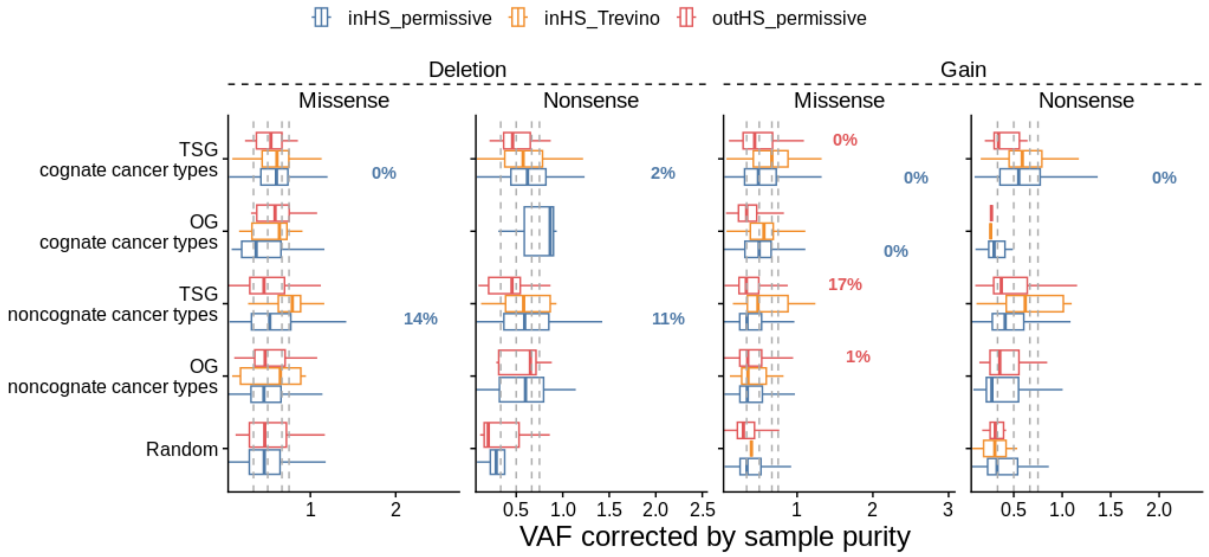


**Supplementary Figure S9. Mutation frequencies in samples with a gene loss or gene gain.** One data point corresponds to one adjusted frequency of one missense or nonsense mutation for each tumor sample from TCGA. FDRs are labelled for FDR ≤ 25%.

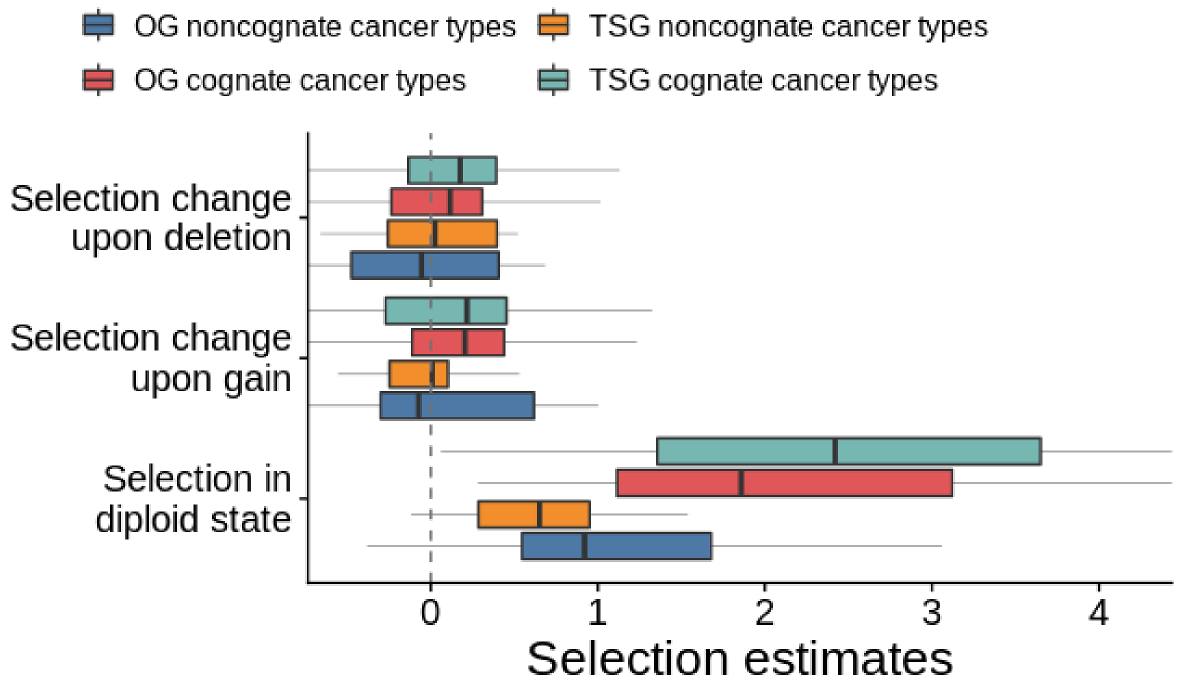


**Supplementary Figure S10. Selection on nonsynonymous mutations in diploid state and conditional selection upon CNA events in GENIE study**. Debiased regression coefficient ω (estimation of selection pressure in the diploid state and debiased regression coefficients δ on the interaction term between the selection variable t and copy number variable c, for gene deletions, and copy gains. One data point corresponds to one gene-tumor type combination.

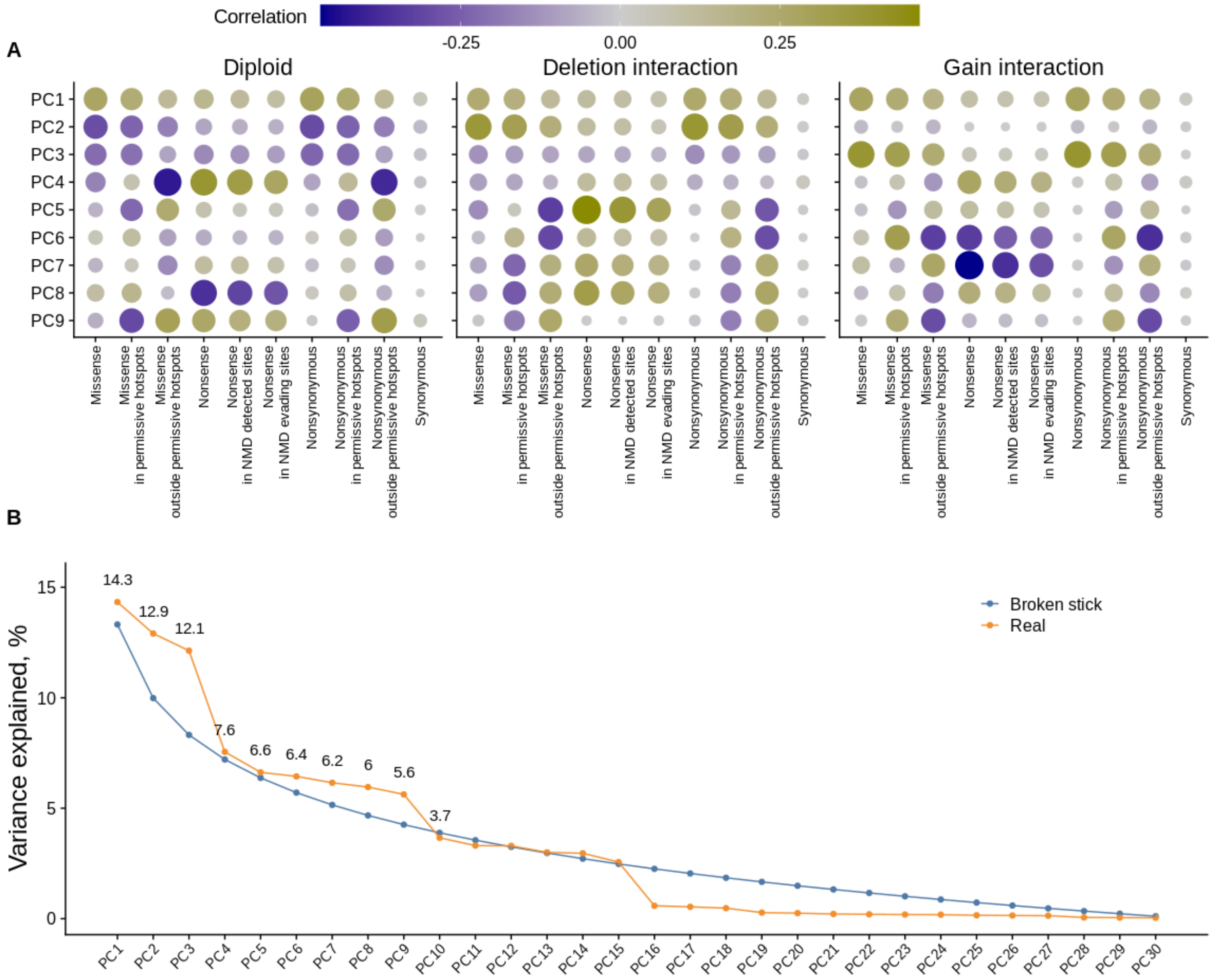


**Supplementary Figure S11. PC analysis of selection in different copy number states. A.** Loadings (correlations with features) of nine first principle components (prior to rotation). **B.** Proportion of variance explained by the derived principle components (orange) compared to the expected explained variance. Three first PCs were found significant (broken stick test).

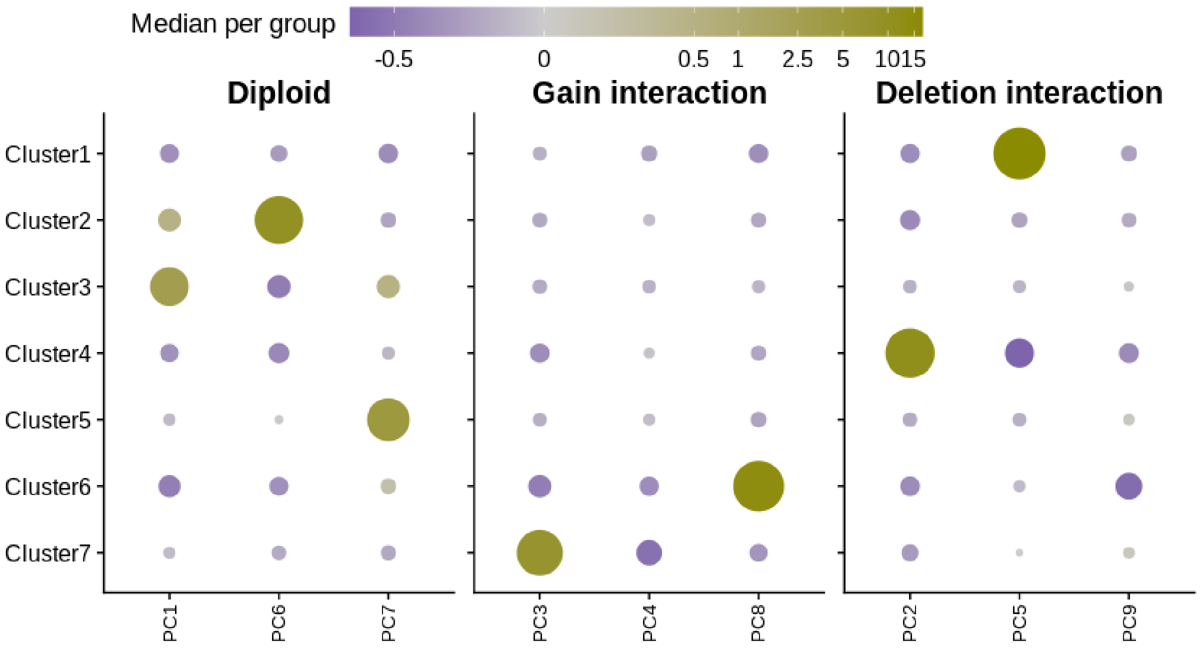


**Supplementary Figure S12. Average factor scores for each cluster.** Factors (rotated PCs) averaged for each cluster of gene-tumor type pairs. Color and size represent the average factor score (scale is logarithmic with a smooth transition to linear scale around 0, which enhances the representation of small values), The size scale is designed to make smaller values more prominent than the color scale, for a more clear visualization.
