## Supplementary Table S1 for "Copy number losses of oncogenes and gains of tumor suppressor genes generate common driver events of human cancer"

**Supplementary Table 1.** Cancer types from the discovery cohort with the largest number of nonsynonymous mutations

| **Cancer** | **Full cancer name** | **Number of mutations** | **Number of samples** |
|---|---|---|---|
| SKCM | Skin cutaneous melanoma | 499 418 | 1004 |
| UCEC-MSI | Uterine corpus endometrial carcinoma MSI tumor samples | 270 215 | 168 |
| LUAD | Lung adenocarcinoma | 212 630 | 996 |
| UCEC-POLE | Uterine corpus endometrial carcinoma hypermutated tumor samples | 205 066 | 47 |
| LUSC | Lung squamous cell carcinoma | 166 235 | 700 |
| COREAD | Colorectal adenocarcinoma | 128 934 | 1121 |
| COREAD-POLE | Colorectal adenocarcinoma hypermutated tumor samples | 124 671 | 27 |
| COREAD-MSI | Colorectal adenocarcinoma MSI tumor samples | 124 388 | 92 |
| BLCA | Bladder urothelial carcinoma | 124 173 | 554 |
| BRCA-Lum | Breast invasive carcinoma luminal subtype | 108 439 | 1340 |
| HNSC | Head and neck squamous cell carcinoma | 99 026 | 762 |
| STAD-MSI | Stomach adenocarcinoma MSI tumor samples | 82 613 | 86 |
| PRAD | Prostate adenocarcinoma | 79 471 | 1530 |
| PAAD | Pancreatic adenocarcinoma | 65 262 | 1011 |
| BRCA | Breast invasive carcinoma | 60 263 | 810 |
| MM | Multiple myeloma | 56 732 | 1089 |
| LGG | Brain lower grade glioma | 53 987 | 588 |
| Kidney | Kidney cancer | 50 870 | 1041 |
| ESAD | Esophageal adenocarcinoma | 50 671 | 463 |
| NSCLC | Non-small cell lung cancer | 49 075 | 117 |
| STAD | Stomach adenocarcinoma | 48 011 | 450 |
| UCEC | Uterine corpus endometrial carcinoma | 46 691 | 454 |
| GBM | Glioblastoma multiforme | 43 163 | 438 |
| PBCA | Pediatric brain cancer | 37 362 | 617 |
| LIHC | Liver hepatocellular carcinoma | 36 784 | 417 |
| CESC | Cervical squamous cell carcinoma and endocervical adenocarcinoma | 36 719 | 334 |
| SKCA | Skin adenocarcinoma | 34 959 | 100 |
| ESCA | Esophageal Cancer | 30 948 | 253 |
| OV | Ovarian cancer | 30 284 | 436 |
| BRCA-tri- | Breast invasive carcinoma triple-negative subtype | 29 983 | 328 |
| LICA | Liver cancer | 26 322 | 193 |
| MALY | Malignant lymphoma | 20 239 | 252 |
| NET | Neuroendocrine cancer | 18 669 | 263 |
| BRCA-HER2 | Breast invasive carcinoma HER2-positive subtype | 16 490 | 16 490 |
